# A generalized observation confirmation model to account for false positive error in species detection-nondetection data

**DOI:** 10.1101/422527

**Authors:** John D. J. Clare, Benjamin Zuckerberg, Philip A. Townsend

## Abstract

Spatially-indexed repeated detection-nondetection data is widely collected by ecologists interested in estimating parameters associated with species distribution, relative abundance, phenology, and more while accounting for imperfect detection. Recent model development has focused on accounting for false positive error as well, given growing recognition that misclassification is common across many sampling protocols. To date, however, the development of model-based solutions to false positive error has been largely restricted to occupancy models. We describe a general form of the observation confirmation protocol originally described for occupancy estimation that permits investigators to flexibly and intuitively extend several models for detection-nondetection data to account for false positive error. Simulation results demonstrate that estimators for relative abundance and arrival time exhibit relative bias greater than 20% under realistic levels of false positive prevalence (e.g., 5% of detections are false positive). Bias increases as true and false positives occur in more distinct places or times, but can also be sensitive to the values of the state variables of interest, sampling design, and sampling efficiency. Results from an empirical study focusing on patterns of gray fox relative abundance across Wisconsin, USA suggest that false positive error can also distort estimated spatial patterns often used to guide decision-making. The extended estimators described within typically improve performance at any level of confirmation, and when false positive error occurs at random and constitutes less than 10% of all detections, the estimators are essentially unbiased when more than 50 observations can be confirmed as true or false positives. The generalized form of the observation-confirmation protocol is a flexible model-based solution to false positive error useful for researchers collecting data with sampling devices like trail or smartphone cameras, acoustic recorders, or other techniques where classifications can be reviewed post-hoc.

## Introduction

Detection non-detection data is widely collected in ecology. The importance of collecting repeated individually-indexed detection-nondetection data to account for observation error associated with imperfect detection has been recognized for decades (e.g., Otis et al. 1978). More recently, spatially-structured models that use repeated observations of species occurrence at specific locations to account for imperfect detection have seen rapid development and increasing use. These models have greatly expanded what can estimated with detection-nondetection data, permitting inference about several ecological state variables such as species distribution, abundance, phenology, density, and associated dynamics (MacKenzie et al. 2002, Royle and Nichols 2003, Roth et al. 2014, Ramsey et al. 2015, Rossman et al. 2016).

However, a growing body of evidence suggests that imperfect detection is not the only important source of observation error in detection-nondetection data. Across a variety of species classification or identification protocols, false positive error (e.g., a species is falsely observed at a time or place when not present) has been shown to interspecifically vary from nearly negligible to constituting 20% of observations or more (Simons et al. 2007, McClintock et al. 2010a, Swanson et al. 2016, Norouzzadeh et al. 2018). Simulation results have shown that even relatively small amounts of false positive error can severely bias species distribution estimators that assume it does not occur (e.g., absolute error in the proportion of occupied sites > 0.1, Miller et al. 2011, Ruiz-Gutiérrez et al. 2016). This has motivated investigators to adopt a variety of strategies aimed at ameliorating biases induced by false positive error. The most thorough strategy is to implement a complete data review after collection (e.g., Gardiner et al. 2012), but this can become burdensome with the sizable datasets produced by automated detection devices or citizen scientists (Ruiz-Gutiérrez et al. 2016). In the interest of efficiency, investigators often turn to several other approaches such as using partial reviews to develop indicators or algorithms to identify misclassified data, simplifying classification tasks, or providing additional training (data) to human (or computer-based) classifiers (Swanson et al. 2016, see review by Kosmala et al. 2016). Although these approaches can greatly improve data quality, they suffer certain limitations. It can be time-consuming to quantify improvements in data quality resulting from protocol or training changes and to determine how accurate raw data or error indicators need to be to achieve desired inferential reliability (Clare et al. 2019). Furthermore, appropriately propagating indicator uncertainty or any remaining data error into subsequent estimate uncertainty is not always straightforward.

Model-based solutions may be the most elegant way to ameliorate false positive error, and several solutions for detection-nondetection data following different protocols have been described specifically for occupancy models (Royle and Link 2006, Miller et al. 2011, Chambert et al. 2015, Ferguson et al. 2015, Ruiz-Gutiérrez et al. 2016). The base model described by Mackenzie et al. (2002) conceptualizes observed absences as a mixture of true and false negatives, and subsequent extensions all conceptualize observed occurrences as also arising from a mixture of true and false positives. The varied protocols primarily differ with regard to how the probability of a false positive observation is informed within the estimation process. Under the ‘full’ estimator described by Royle and Link (2006), all observed occurrences are of unknown reliability, and disentangling the false negative and false positive mixtures requires multiple constraints. Subsequent developments leverage auxiliary data to directly inform parts of the positive and/or negative mixtures, like additional error-free observations provided by an additional sampling method (site-confirmation protocol), experimental trials for classification performance under controlled settings (calibration protocol) or confirming a subset of the observations post-hoc (observation confirmation protocol; Miller et al. 2011, Chambert et al. 2015, Ferguson et al. 2015, Ruiz-Gutiérrez et al. 2016). An advantage of these model-based solutions is that they reduce positive estimator bias associated with false positives while also propagating uncertainty in observation reliability into estimator uncertainty.

The rapid development of model extensions to accommodate false positive error across a variety of sampling situations suggests that many researchers recognize that false positives are common and problematic. It is surprising and concerning, then, that model development and application has largely been restricted to occupancy estimation. False positives presumably induce bias in any parameter or state variable estimated with detection-nondetection data, and the cost of biased estimation is arguable greater for many parameters or state variables other than distribution (e.g. species abundance).

We suspect that a major barrier to broader adoption and development is that many of the specific conditional probability statements described within false positive occupancy models are not intuitive for many ecologists to translate into other models using similar data. Here, we clarify the flexibility of model-based solutions to false positive error within detection-nondetection data using basic laws of probability. We focus on the observation-confirmation protocol, which is most pragmatic formulation for investigators reliant upon sampling techniques that produce data that can be reviewed *a posteriori* (images, audio recordings, physical specimens like scats, etc.) and is arguably the most efficient model-based solution (Chambert et al. 2015). Here, we present a generalizable model structure, demonstrate that standard and false positive occupancy models are special cases of said structure, and present extensions of models

### The Generalized Model Form

Throughout, we assume a data matrix of binary observations ***y*** that corresponds to the detection or nondetection of some species (or other phenomena of interest) at *i =* 1, 2, … R locations over *j* = 1, 2…T discrete sampling intervals. We also assume *y_i,j_*~ Bernoulli (*θ*), where *y_i,j_* = 1 if a species is observed and 0 otherwise; *θ* describes the probability of a positive observation. Commonly, *θ* is likely to be a mixture of the respective probabilities of true positive observations that are the research focus (*θ_tp_*) and false positive observations (*θ_fp_*); decomposing these components allows one to make exclusive inference about *θ_tp_*. Assuming that true and false positive observations are independent, the specification of the mixture of true and false positives can be derived from addition rules for probabilities. True and false positives might be mutually exclusive events such that *y_i,j_* ~ Bernoulli (*θ_tp_* + *θ_fp_*), implying that a positive observation is either a true positive or a true positive. Alternatively, a specific binary outcome might include only true positive observations, only false positive observations, or both, in which case *y_i,j_* ~ Bernoulli (*θ_tp_*×[1– *θ_fp_*] + *θ_fp_*× [1– *θ_tp_*] + *θ_tp_*×*θ_fp_*). After factorization, this simplifies to *y_i,j_* ~ Bernoulli (*θ_tp_* + *θ_fp_*– [*θ_tp_*×*θ_fp_*]). These probabilistic statements constitute a generalized structure for modeling true and false positive probabilities in binary data.

Many models commonly used by ecologists can be described using the generalized structure. A generalized (logit) linear model for species occurrence could be described as a case that assumes logit 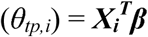 and *θ_fp_* = 0. A generalized (logit) linear mixed model for repeated species occurrences is a case assuming logit 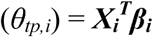, *θ_fp_* = 0, and *β****_i_*** ~ Normal(***µ****_β_*, σ*_β_*). The occupancy model described by MacKenzie et al. (2002) can be described as a case assuming *θ_fp_* = 0 and *θ_tp,i_* = *z_i_* × *p*, where *z_i_* is the latent occupancy state of site *i, z_i_* ~Bernoulli (*ψ*), *p* is the probability of detecting an organism at a site given that it is present, and both *p* and *ψ* might be assumed to vary following other functions. Importantly in these cases, if *θ_fp_* > 0, *θ_tp,i_* is estimated incorrectly. Rather, the estimands associated with *θ_tp,i_* actually describe a mixture of true and false positive probabilities, either *θ_tp,i_*+ *θ_fp_*, or *θ_tp_* + *θ_fp_* – (*θ_tp_*×*θ_fp_*).

Model-based solutions for false positive error in occupancy models explicitly describe observations using the mixture specifications for *θ_tp_* and *θ_f**p**_* described above. The full, site-confirmation, and calibration models follow the mutually exclusive specification, where *y_i,j_*~ Bernoulli (*θ_tp,i_*+ *θ_fp_*). Ignoring the auxiliary components of the site-confirmation and calibration models, all three generally treat *θ_fp,i_* = (1 – *z_i_*) × *p_10_* and *θ_tp,i_*= *z_i_* × *p_11_*, where *p_11_* is the probability of detecting an organism during an interval at a location where it is present, and *p_10_* is the probability of detecting an organism during an interval at a location where it is not present. In fact, because *θ_fp,i_* and *θ_tp,i_* are dependent upon the state *z_i_*, true and false positive detections are mutually exclusive not only within specific cells *y_i,j_*, but across sites (rows) ***y_i,-_***. The observation-confirmation model does not require that true and false positives be mutually exclusive, and we focus on extending this model below.

### The Observation Confirmation Protocol

As originally described by Chambert et al. (2015), the observation confirmation model addresses an estimation problem where the purpose is to estimate the binary occurrence state within a finite sample of sites (*z_i_*) or the population level probability of occurrence *ψ*, assuming *z_i_* ~ Bernoulli (*ψ*). Within some subset of sampling intervals *j* at specific sites, some observations (*v_i,j_*) have been verified as either containing only true positives (*v_i,j_*= 0), only false positives (*v_i,j_* = 1), both (*v_i,j_*= 2), or no positive observations (*v_i,j_* = 3); note that we order the terms differently than Chambert et al. (2015). These verified observations might include physical samples (hair or scat), images, audio or video recordings, or any other data type that can subsequently be confirmed via expert evaluation, laboratory analysis, or other means. The confirmed samples *v_i,j_* ~ Categorical (Ω), where the elements of Ω are conditional on whether the species occurs at site *i* or not. If *z_i_*= 1 (species occurs), then Ω**_i_** = [{(*s_1_*× (1 – *s_0_*)} {*s_0_*× (1 – *s_1_*)} {*s_1_*× *s_0_*} {(1 – *s_1_*) × (1 – *s_0_*)}]. If *z_i_*= 0, the only possible outcomes are a false positive detection or no detection, and Ω**_i_** = [{0} {*s_0_*} {0} {1 – *s_0_*}]. Here *s_1_* and *s_0_* represent the respective probabilities that a sampling interval contains > 0 true positive or false positive observations conditional upon the occurrence state. The similar conditional statement for unconfirmed data is *y_i,j_*~ Bernoulli (*z_i_* × *p_11_* + (1 – *z_i_*) × *p_10_*). If present, a species can either be truly or falsely detected (*y_i,j_* = 1| *z_i_*= 1) with probability *p_11_* = *s_1_* + *s_0_*– (*s_1_* × *s_0_*), and if not (*y_i,j_*= 1| *z_i_* = 0), it can only be falsely detected with probability *p_10_* = *s_0_*.

The observation confirmation occupancy model thus follows a non-exclusive form of the general model based upon addition rules for probabilities. The first three elements of Ω given that *z_i_* = 1 in mirror the constituent terms in the general model *θ_tp_*(1– *θ_fp_*) + *θ_fp_*(1– *θ_tp_*) + *θ_fp_θ_tp_*, and the statement for *p_11_* mirrors the factorized *θ_tp_* + *θ_fp_*– (*θ_tp_θ_fp_*). In fact, the observation confirmation model can be described as a specific case of the generalized structure in which *θ_tp,i_* = *z_i_*× *s_1_* and *θ_fp_* = s_0._ The row vector Ω**_i_** is then specified as = [{*θ_tp,i_*× (1– *θ_fp_*)} { *θ_fp_*× (1– *θ_tp,i_*)} { *θ_fp_*×*θ_tp,i_*} {(1 – *θ_tp,i_*) × (1 – *θ_fp_*)}]. As before, *v_i,j_* ~ Categorical (Ω**_i_**). In turn, *y_i,j_* ~ Bernoulli (1 – Ω_4,*i*_); which is exactly equivalent to describing *y_i,j_* ~ Bernoulli (*θ_tp,i_*+ *θ_fp_* – [*θ_tp,i_*×*θ_fp_*]). In full, a hierarchical description of the complete data likelihood is:

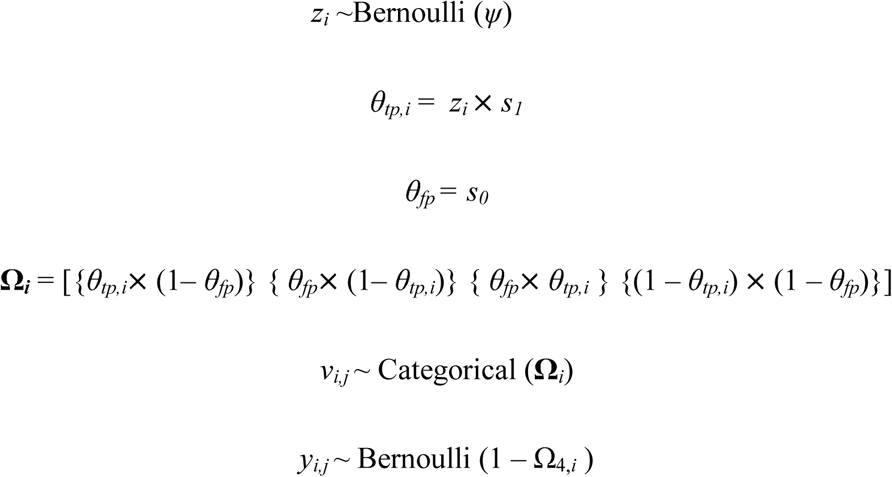

### Extending the Observation Confirmation Protocol to other Models

One outcome of defining the observation confirmation protocol using the general model structure is that it clarifies extension to other models reliant upon detection-nondetection data. One simply needs to update the definition of *θ_tp_* to reflect the full probabilistic statement for a positive outcome using whatever particular model the investigator is interested in fitting (Figure 1). We describe two example extensions in the main text.

**Figure 1.**
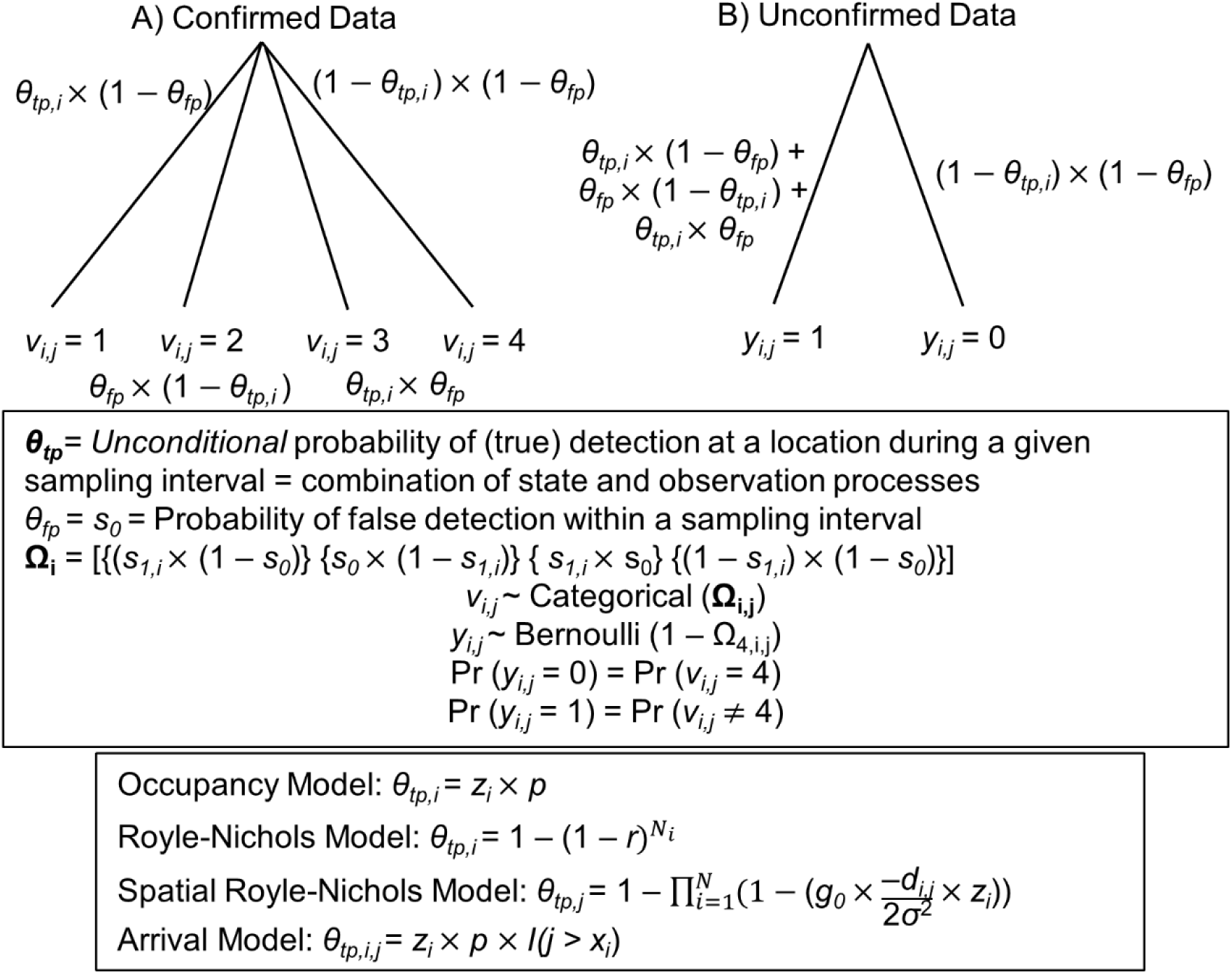
Schematic for the generalized observation-confirmation model for false positive error presented here. Unconfirmed outcomes are positive if the species has either been truly or falsely detected, or negative if neither true nor false detection has occurred. Confirmed data can either result in a true detection only, a false detection only, both, or no detection. The probability of > 0 false detections within a verified sampling interval is denoted *θ_fp_*, while the probability of > 0 true detections, *θ_tp,i_*, is treated as a combination of functions associated with state and conditional observation models.

### Royle-Nichols Model

Royle and Nichols (2003, RN hereafter) describe the unconditional probability of detection (i.e., *θ_tp,i_*) as 1 – (1 – *r*)*^N_i_^*, where *r* is the probability of detecting an individual during a sampling interval and *N_i_* ~ Poisson (λ) denotes the abundance of a species at site *i*. The hierarchical likelihood for a version incorporating false positives following the observation-confirmation protocol is then:

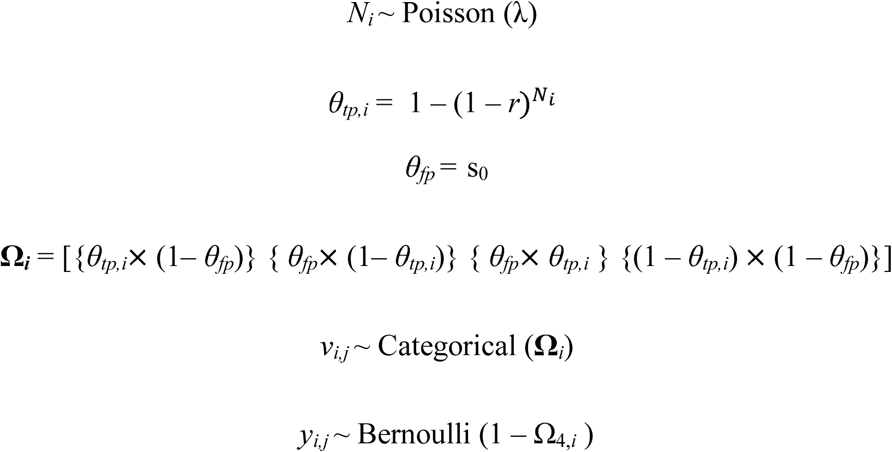

### Phenological ‘arrival’ model

Incorporating false positives following the observation-confirmation protocol within an occupancy model designed to estimate the timing of some ephemeral phenomena such as migration arrival or emergence from torpor (Roth et al. 2014, arrival model hereafter) follows Chambert et al.’s (2015) description except that organisms can only be truly detected during sampling intervals at occupied sites after arrival. Thus, *θ****_tp_*** must be described using indexing for *i* locations and *j* time periods (i.e., *θ_tp,i,j_*). Let arrival time at site *i* be denoted as *x_i_* and assume that *x_i_* ~ Poisson (*φ*). To simplify presentation, we define *x_i_* in terms of sampling intervals *j* rather than specific dates. The hierarchical likelihood is:

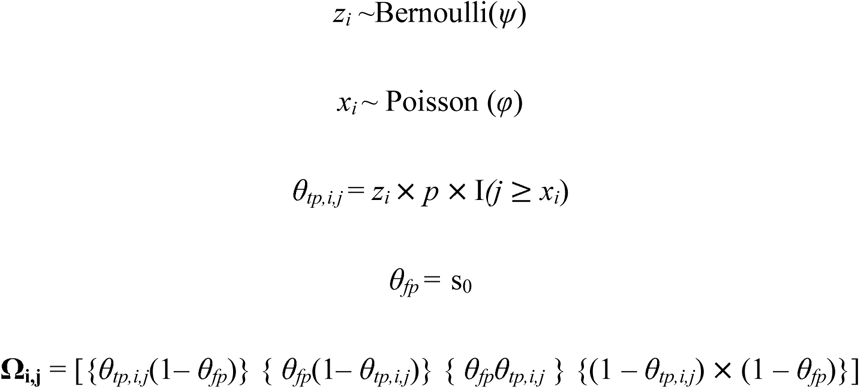

Here, I(*j ≥ x_i_*) is an indicator function denoting whether the species has arrived at site *i*. As before, confirmed and unconfirmed observations can be described as *v_i,j_* ~ Categorical(Ω**_i,j_**) and *y_i,j_* ~ Bernoulli (1 – Ω_4,i,j_).

Only a small sample of the possible extensions are described above. Any number of dynamic or integrated models that incorporate repeated detection-nondetection data can be extended to accommodate false positive error following the observation-confirmation protocol simply by defining *θ****_tp_*** using the model described in the initial paper.

In principle, it is also possible to describe false positive extensions to the RN or arrival models following the site-confirmation or calibration protocol and the full estimator. The site-confirmation and calibration models require that *θ_fp_* be constrained such that for the RN model, false positives are only possible at sites where *N_i_* = 0, and for the arrival model, false positives are only possible at occupied sites during time periods prior to arrival or at unoccupied sites. The general way to extend these models is to define *θ****_tp_*** following the initial description of the model of interest, and to condition *θ_fp_* = 0 when *θ_tp_*_,i_ > 0 and > 0 when *θ_tp_*_,i_ = 0 (see Appendix S1). However, these protocols depend upon *θ_tp_*_,i_ = 0 having functional support. An example where *θ_tp_*_,i_ = 0 is impossible and where extension is limited to the observation-confirmation protocol is the spatially explicit variant of Royle and Nichols’ (2003) model (Ramsey et al. 2015; see description in Appendix S1).

### Exploring Base Model Sensitivity to Error and Performance of Extended Models

We undertook a proof of concept simulation study to evaluate the baseline sensitivity of the RN and arrival models to different amounts of false positive error and the performance of the described extensions. Throughout, we considered a fixed level of effort—200 spatial replicates with 20 temporal replicates each. Given the wide range of potential variability in parameter values, functional forms, and sensitivity to observation aggregation (Balantic and Donovan 2019), this is not an exhaustive assessment of estimator properties.

### Royle-Nichols Model

We first considered 6 different simulation scenarios, within which we generated 300 replicate datasets with site-specific abundances N*_i,sim_* ~ Poisson (λ*_i,sim_*) and log (λ*_i,sim_*) = *β_0_+ β_1_X_1,i,sim_*, where *X_1,i,sim_* ~ N (0, 1), *β_0_* = 0 or −1.5 (3 scenarios each), and *β_1_* = 1; *sim* indexes a particular simulation replicate. Thus, at a site with an average simulated covariate (*X_1_*_,i_ = 0), expected abundance was respectively 1 animal or roughly 0.25 animals. These values were chosen because the RN model tends to perform best when site-specific abundance is low (Kéry and Royle 2016, p. 302) and is perhaps most commonly applied for low-density species. We first generated ‘true’ detection data as Bernoulli (*p_i,sim_*), where *p_i,sim_* = 1 – (1 – *r_i,sim_*)^Ni,sim^, logit (*r_i,sim_*) = *α_0_* + *α_1_X_2,i,sim_*, *X_2,i,sim_*~ N (0, 1), *α*_0_ = –1.73, and *α*_1_ = 1. Thus, an individual at an average site was expected to be detected with a probability of about 0.15 per sampling occasion.

Within these scenarios, we generated false-positive detections as occurring at random across all site intervals within a simulation. The probability of a false-positive detection within a cell was derived such that *θ_fp_*(*s_0_*) constituted approximately 1%, 5% or 10% of *θ_tp_*+ *θ_fp_* with random Binomial variance (absolute values of *s_0sim_* ranged from < 0.001 to roughly 0.025). We defined *θ_fp_* proportionally here rather than explicitly exploring specific values for the parameter itself because few studies provide empirical estimates of false positive probabilities, and many report percentages of observations that are true or false positives (e.g., Simons et al. 2007, Swanson et al. 2016, Norouzzadeh et al. 2018). Furthermore, proportional definitions are also commonly used to define thresholds for accurate data (e.g., 90% accuracy or 95% accuracy; McShea et al. 2016, Swanson et al. 2016).

Within each scenario, we sampled positive detections at random to create the verified data *v_i,j,sim_* (the number of verified samples = {10, 20, 30, 40, 50, 60, 70, 80, 90 100} with 30 replicates for each level per scenario). Each of the 1800 generated datasets was used to fit both a standard RN model and a false-positive extension. We evaluated the performance of both estimators (relative bias, root squared error, standard deviation of the posterior distribution, coefficient of variation, and frequentist coverage—% of 95% CI that included the true value) with respect to *β*, *α*, and the finite sample population size (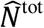, derived as 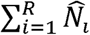).

The initial scenarios were primarily selected to evaluate estimator performance along previously demonstrated gradients of sensitivity or precision: we expected the base estimator to exhibit more bias as the ratio of *θ_fp_* to *θ_tp_* increased (Clare et al. 2019), and expected the extended model to exhibit less relative bias, and estimate uncertainty with more verified samples and a smaller proportion of false positives (Miller et al. 2011, Chambert et al. 2018). In practice, the generating process for false positive error is probably rarely constant given that misclassification probably arises from confusion between phenomena that are not randomly distributed (e.g., species). Misclassification in detection-nondetection data is essentially spatial or spatiotemporal error, and estimator sensitivity may more precisely depend upon similarity in the location or time of true and false positive observations. To demonstrate this, we considered two subsequent scenarios where β_0_ = −1.5 and β_1_ = 1 with *α* defined as before, and logit(*s_0,i_*) = −6 +{−1, 1}*X_1,i,sim_* such that false positive observations were either negatively or positively associated with abundance and the probability of true positive observations. The verification protocol was simulated as previously. Empirically, simulated false positive observations constituted about 6% of all observations, within the range of values considered previously. We fit 3 models for each of these scenarios: one in which *s_0_* was assumed = 0, one in which it was assumed to be constant, and one in which it was modeled as varying in relation to *X_1,i,sim_*.

### Arrival Model

Our exploration of the arrival model was similar. We first considered three scenarios with the following parameterization: logit (*ψ_i,sim_*) = *β_0_+ β_1_X_1,i,sim_, X_1,i,sim_*~ N (0, 1), *β_0_* = 0, and *β_1_* = 0.5; logit (*p_i,sim_*) = *α_0_ + α_1_X_2,i,sim_, X_2,i,sim_* ~ N (0, 1), *α_0_* = –2, *α_1_* = 0.5, and average arrival time *φ* = occasion 6. True observations *y_i,j,sim_* were generated as Bernoulli (*z_i,sim_* × *p_i,sim_* × I(*a_i,j,sim_*)), where *z_i,sim_* ~ Bernoulli (ψ_i,sim_), and site and simulation specific arrival time *a_i,j,sim_* ~ Poisson (*φ*). That is, the occupancy probability of an average site was 0.50 and the probability that of true detection conditional on arrival and occupancy at an average site was *ca*. 0.12. We simulated 300 replicates per scenario; as before, false positive detections varied as *s_0sim_* ≅ {0.01, 0.05, 0.10} × *y_sim_* across scenarios, and the size of *v_i,j,sim_* ranged from 10-100. We fit both the standard arrival model and the false-positive extension to each simulation replicate.

We next considered 6 additional scenarios to explore other sources of sensitivity. We defined logit (*s_0,i,j,sim_*) = −6 +{–1, 1}*X_1,i,sim_* for *j* > 4 and *s_0,i,j,sim_* =0 for *j* ≤ 4, and *α*_1_ = {–2, –1} (at an average site, *p ca.* 0.27 on the real scale) for four scenarios, and logit (*s_0,i,j,sim_*) = −6 +{–1, 1}*X_1,i,sim_* for *j* > 2 and *s_0,i,j,sim_*= 0 for *j* ≤ 2, and *α*_1_ = –2 for two others; other values followed the previous description. While the previous three simulation scenarios considered allowed false positives to happen at random, the formulation for these might be more realistic if, for example, false positives are mostly associated to look-alike species that arrive just slightly or somewhat earlier than the focal species and have either comparable or distinct habitat associations. Empirically, these different formulations for false positive error resulted in false positives accounting for roughly 6% (*α*_1_ = −1 and *s_0,i,j,sim_* = 0 for *j* ≤ 4), 7% (*α*_1_ = −1 and *s_0,i,j,sim_* = 0 for *j* ≤ 2), and 3% (*α*_1_ = −2 and *s_0,i,j,sim_* =0 for *j* ≤ 4) of all observations. As with the RN model, we fit models that assumed *s_0_* = 0, models that assumed *s_0_* was a constant, and models that (correctly) assumed *s_0_* varied in relation to X_1,i,sim._ We evaluated estimator properties with respect to *α*, *β*, 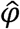 and finite sample estimate of the proportion of occupied sites (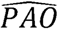, derived for each simulation as 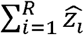).

### Evaluating transferability of s_0_

A potentially appealing property of the generalized description is that the confirmation outcomes that primarily contribute to the estimation of *θ_fp_* and the parameter’s underlying generating process are likely to be consistent regardless of how *θ_tp_* is formulated. This suggests that if data or computational resources are lacking, one might be able to use an informative prior for *θ_fp_* given previous estimates of the parameter from a distinct (and more quickly fit) model. To briefly explore transferability, we fit the original observation confirmation occupancy model described by Chambert et al. (2015; i.e., *θ_tp,i_* = *z_i_×p_i_*) to every simulated dataset described previously, and then fit a model with the correct (i.e., RN or arrival) structure for *θ_tp,i_* and for which *θ_fp_* was strictly informed by a prior distribution derived from the posterior distribution of the false positive parameter estimated by the occupancy model.

### Application: Predicting Gray Fox Relative Abundance across Wisconsin

The goal of the simulation study was primarily to evaluate the statistical properties of the standard and extended estimators. However, models for detection-nondetection data are also often used to predict or map in order to spatially delineate or prioritize management actions (e.g., Guélat and Kéry 2018). As a case study, we focus upon the relative abundance of gray fox (*Urocyon cinerargenteus*) in Wisconsin, USA. The species has not been recently and systematically monitored or surveyed across the state, and its distribution is poorly understood. Here, we use data from the monitoring program Snapshot Wisconsin (Clare et al. 2019), in which trail cameras are deployed by citizen scientists and classified via a crowdsourcing platform, to investigate spatial patterns in fox relative abundance using the RN model. Gray foxes were known to be frequently misclassified within the dataset (Clare et al. 2019 estimated the probability of a single image being a false positive as 25%; 95% CRI = 18% – 33%). We modeled variation in fox expected abundance as a function of 5 environmental covariates and a spatial smoothing term, with additional covariates used to explain variation in the detection probability of individual foxes and the probability of false positive detections (details in Appendix S3). We used indicator variable selection (Kuo and Malick 1998) to identify important predictors and regularize the log-linear coefficients; we make statewide predictions by applying model-averaged coefficients across a 2 × 2 km lattice.

We fit all models using JAGS v 4.0 (Plummer 2003) to perform Markov-Chain Monte Carlo simulation through R v 3.4 (R Core Team 2017). Appendix S3 and Data S5 provide additional details and code associated with the case study, and Appendix S4 provides code used to implement the simulation study. Throughout the simulation study, we did not simulate any confirmation of non-detections (i.e., simulated confirmations were randomly drawn from positive observations) following Clare et al. (2019). This is because it seems inherently inefficient for practitioners to wade through potentially many images or recordings of non-target species within a putative period of non-detection: practically speaking, confirming ‘absence’ is not always easy to do. Provided the likelihood for *v* still incorporates some probability for non-detection (even if this event is never observed), there seems to be no cost to doing so (Clare et al. 2019).

## Results

As expected, when false positive error was essentially random, standard estimators for abundance, proportion of area occupied, and arrival time were biased, and more biased when false positives constituted a greater proportion of detections (Figures 2 and 3; more comprehensive results in Appendix S2). Random misclassification across all time periods and locations constituting 1, 5 or 10% or all detections led to respective relative biases of 10, 40 and 70% for abundance (across both simulated expected abundances of both 0.23 and 1 individual per site), 3, 10 and 20% for proportion of area occupied, and 3, 20, and 40% for arrival time. The extended estimators exhibited less bias and root mean squared error than the standard estimators regardless of the size of the verified sample; estimator performance with regard to these metrics asymptotically improved as more samples were verified, with minimal marginal improvement once 25 sampling occasions were confirmed (Figures 2 and 3). The extended estimators for species distribution and abundance also exhibited greater uncertainty than the standard estimators, although this difference similarly shrank as more samples were verified. Uncertainty patterns were reversed for arrival time (Figure 3), suggesting that false positive error was inducing overdispersion in arrival relative to Poisson expectations.

**Figure 2.**
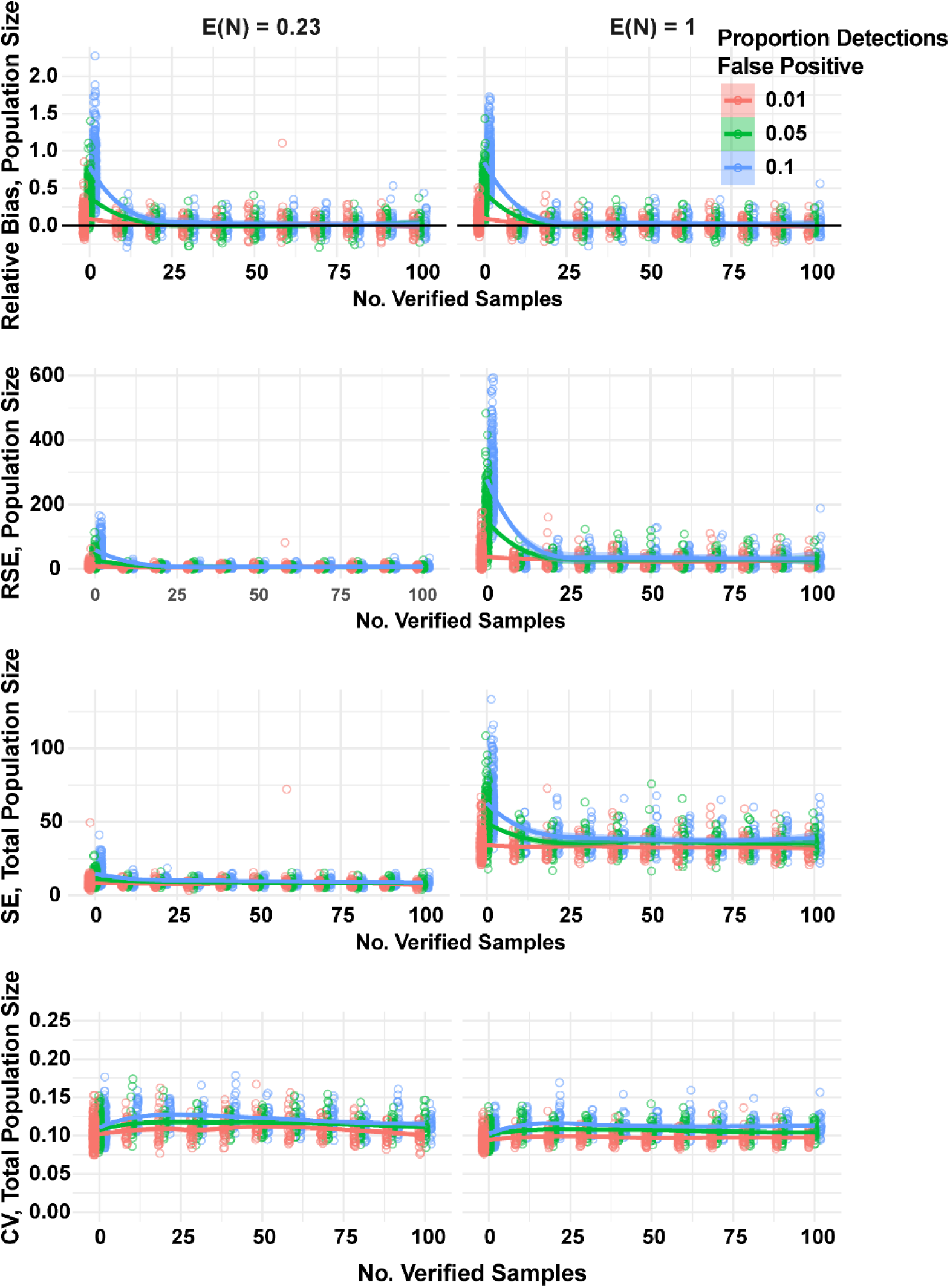
Performance of the standard Royle-Nichols model (when 0 observations are confirmed) and the model extension for false positive error with regard to finite-sample population size under varying levels of random false positive error (% of total observations = 1, 5, or 10) and confirmation effort. SE = standard deviation of the posterior distribution, CV = coefficient of variation.

**Figure 3.**
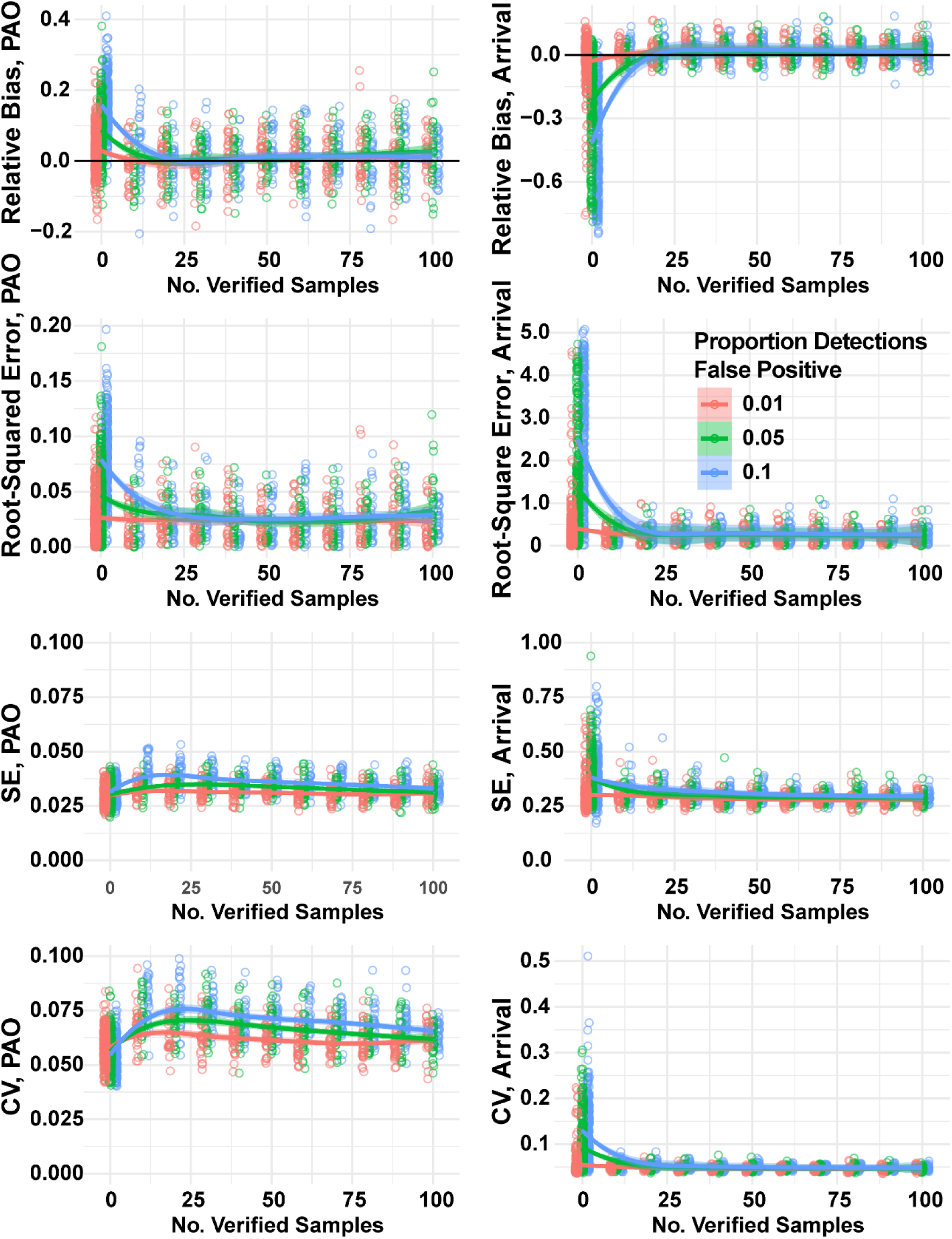
Performance of a standard phenological occupancy model (when 0 observations are confirmed) and the model extension for false positive error with regard to proportion of area occupied and the time of arrival under varying levels of random false positive error (% of total observations = 1, 5, or 10) and confirmation effort. SE = standard deviation of the posterior distribution, CV = coefficient of variation.

### Covariance between θ_fp_ and θ_tp_

The differences became more nuanced when true and false positives covaried in different ways. Estimation of relative abundance, proportion of area occupied, and arrival time using the extended models was unbiased or nearly unbiased regardless of how false positive error was specified (less than 5% relative bias, Figures 4, 5, and 6; Appendix S2). As with random error, extended estimator performance asymptotically improved with a larger confirmation sample. The standard estimators for population size and proportion of area occupied were more biased when false positive detections were more likely at locations where true positive detections were less likely (i.e., when false positive probability was negatively associated with a covariate itself positively associated with expected abundance or occupancy, Figures 4 and 5). Unsurprisingly, estimates of arrival time using the standard estimator were insensitive to spatial patterns in error, but were more biased as the simulated initiation of false positives occurred earlier relative to the average true arrival time (Figure 6). In fact, when false positives were simulated as starting only one sampling occasion before true positives, the standard estimator exhibited very slightly less bias and RMSE than the generating estimator (indeed, the extended estimator always exhibited some slight positive bias with regard to arrival time). This appeared to be a form of small sample bias associated with low simulated detection probability, as the standard model fit to the ‘true positive’ simulated data was equivalent to the extended model (Appendix S2). For the standard model fit to data with simulated error, this appears to be a specific realization of offsetting biases (low detection probability offsetting some minimal difference in the start times of false and true positives).

**Figure 4.**
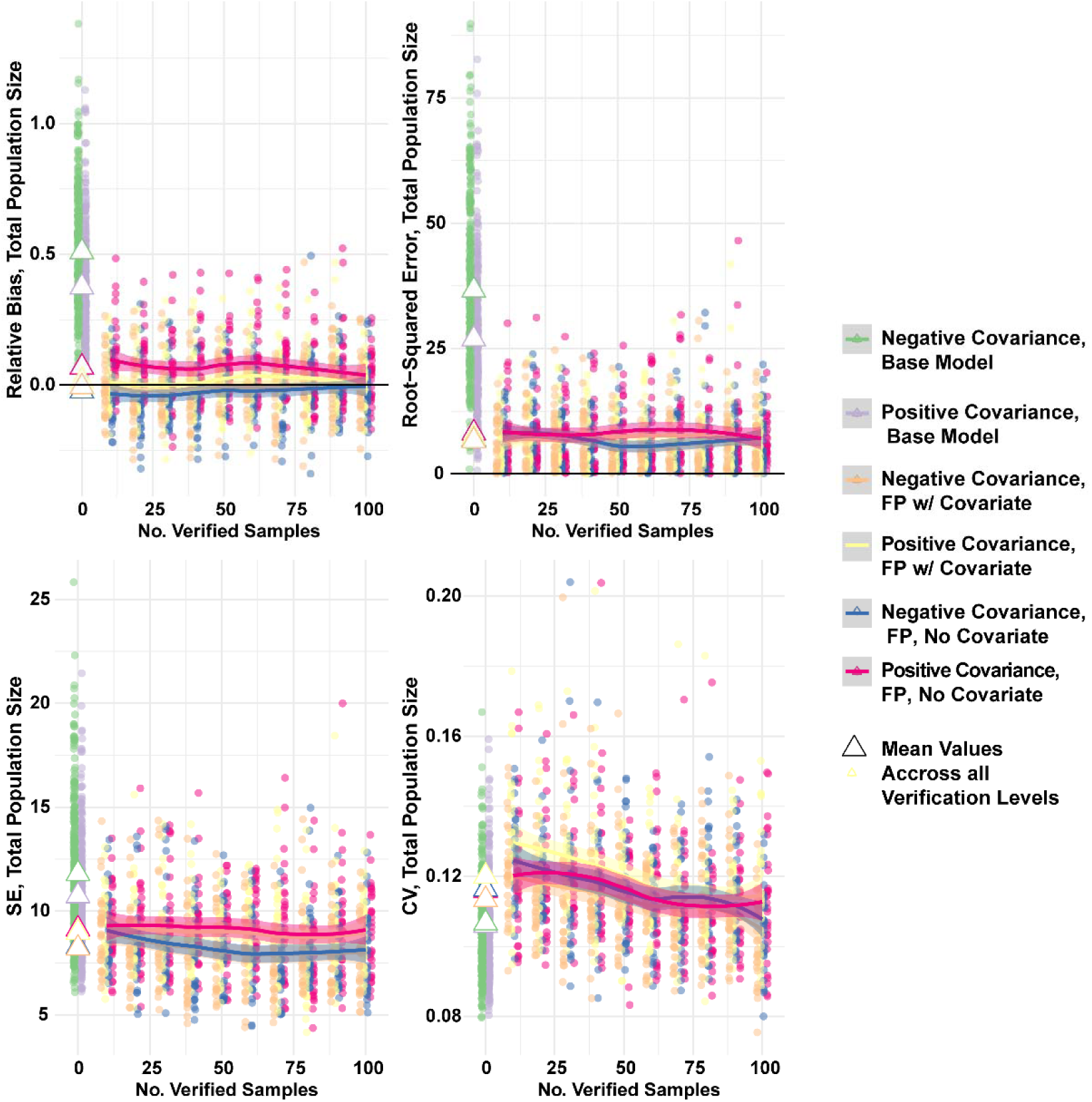
Performance of the standard Royle-Nichols model (when 0 observations are confirmed) and the model extension for false positive error with regard to finite-sample population size under different directional covariance between true and false positive detections, and when different models for false positive error (either a misspecified constant model or the generating model) were fit. SE = standard deviation of the posterior distribution, CV = coefficient of variation.

**Figure 5.**
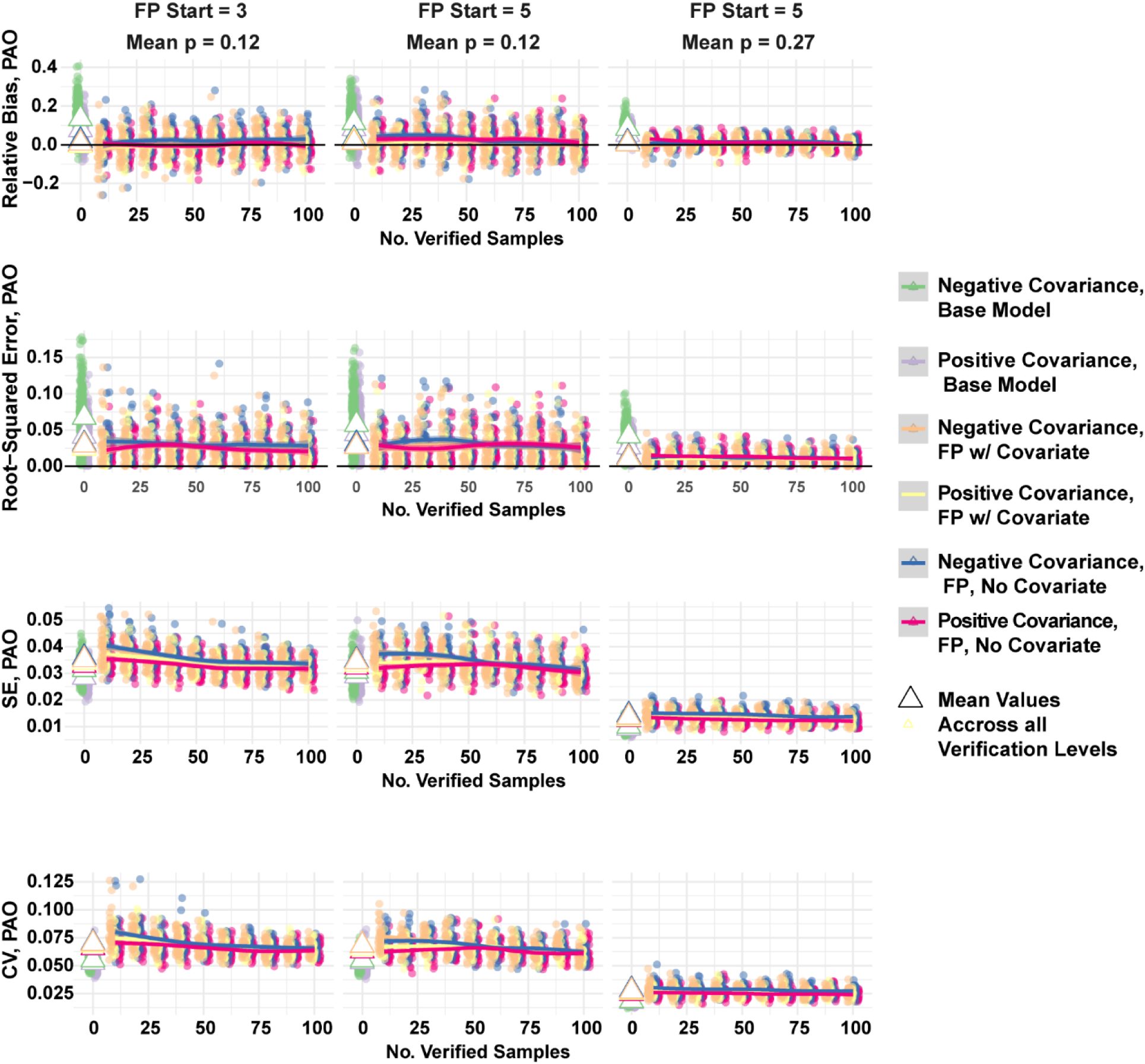
Performance of the standard standard phenological occupancy model (when 0 observations are confirmed) and the model extension for false positive error with finite-sample population size with regard to proportion of area occupied under different directional covariance between true and false positive detections, and when different models for false positive error (either a misspecified constant model or the generating model) were fit. SE = standard deviation of the posterior distribution, CV = coefficient of variation.

**Figure 6.**
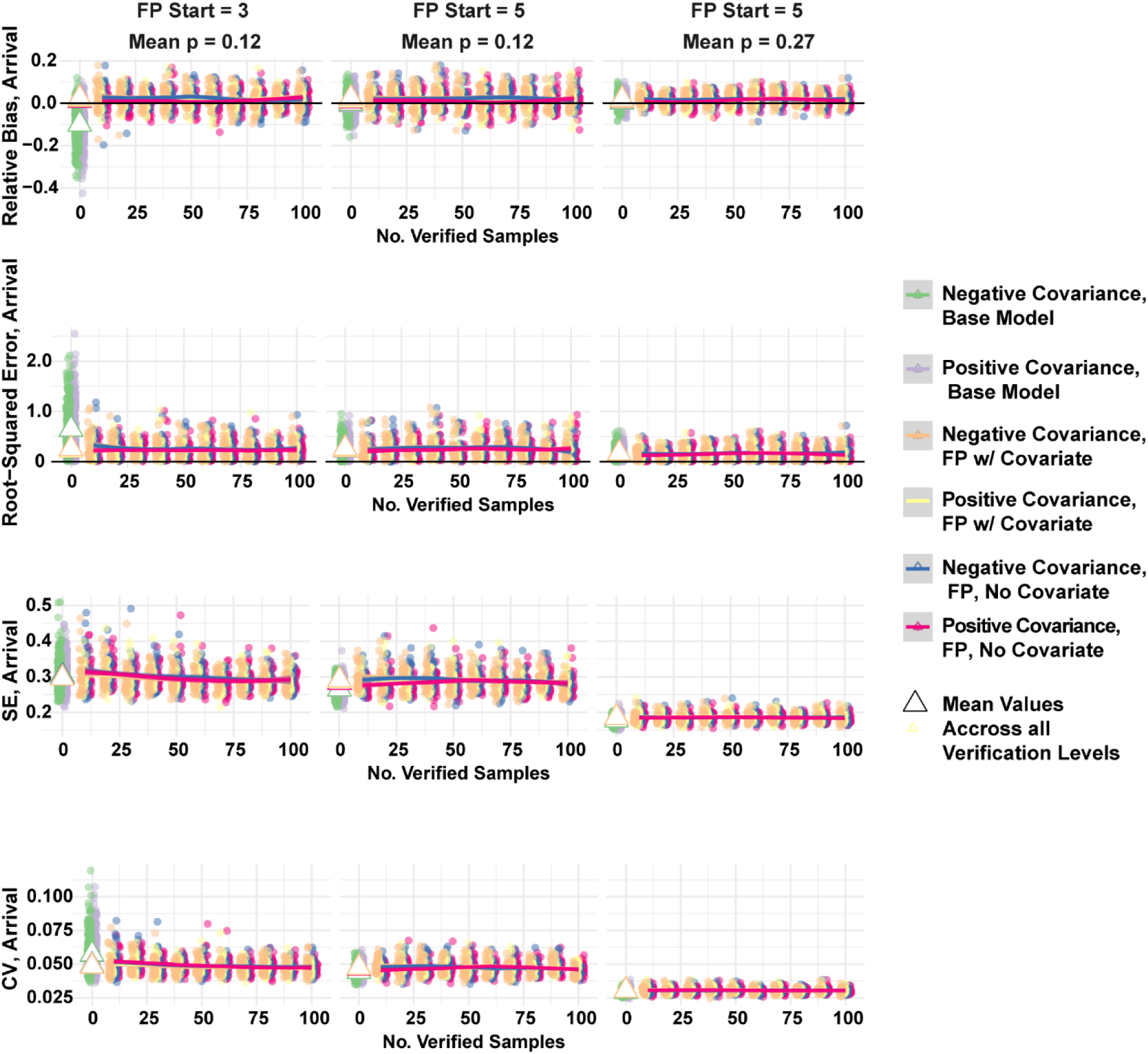
Performance of the standard standard phenological occupancy model (when 0 observations are confirmed) and the model extension for false positive error with finite-sample population size with regard to estimated arrival time under different directional covariance between true and false positive detections, and when different models for false positive error (either a misspecified constant model or the generating model were fit). SE = standard deviation of the posterior distribution, CV = coefficient of variation.

In contrast, estimation of the proportion of area occupied became more biased as simulated false positive error started earlier (Figure 5). Moreover, occupancy estimation became more biased with when the conditional probability of truly detecting a species was lower, further indicating some potential sensitivity to small-sample bias. Arrival estimation was primarily sensitive to the conditional probability of detection insomuch as estimate uncertainty increased (Figure 6).

### Transferability of s_0_

Using an informed prior for false positive error generally resulted in parameter estimates that were strongly correlated with estimates produced when confirmation results were directly incorporated into the likelihood, particularly for the RN model (Appendix S2, Figure S1). There was more variability between point estimates of proportion of area occupied and time of arrival derived from models using an informed prior vs. incorporating the confirmation data. This did not appear to be related to the size of the verification sample; instead, relative to including confirmed observations in the likelihood, using an informed prior tended to result in slightly smaller estimates of proportion of area occupied and slightly larger estimates of arrival time (Appendix S2, Figure S2). Overall, estimator performance scarcely differed when using an informed prior rather than including confirmed locations (Appendix S2, Tables S1 – S9).

### Case Study

Within our review of putative gray fox images, 67% were correctly classified; after aggregation within 179 distinct sampling occasions, 60% consisted of only true positives, and 40% consisted of only false positives (either coyote, *Canis latrans* or red fox, *Vulpes vulpes*). Both the standard and extended models estimated a negative association between fox abundance and the proportion of surrounding coniferous forest. Indicator variable selection provided less support for the inclusion of all covariates within the standard model than the extension (Appendix S3, Tables 1 and 2), and although patterns in predicted gray fox abundance were similar overall (r = 0.80 for pixel-wise point estimates, Figure 7), predictions from the standard model more strongly depict a purely spatial tend with less texture and pronounced discrepancies within the western part of Wisconsin. Although not a primary objective of this modeling exercise, we note the point estimate for expected state-wide population size using the standard model was more than 300% greater than when false positives were accounted for, although estimates overlapped substantially due to imprecision induced by the spatial terms (posterior median = 14999, 95% CRI = 1982 – 255660, 80% CRI = 4754 – 34197 for the model assuming only false negatives, vs. 4773, 95% CRI = 205 – 215225, 80% CRI = 337 – 21341; Figure 7). These differences manifested despite a trivially small estimate of the probability of a false positive detection per sampling interval (at an average site, 0.15%, 95% CRI 0.12% – 0.18%, Appendix S3).

**Figure 7.**
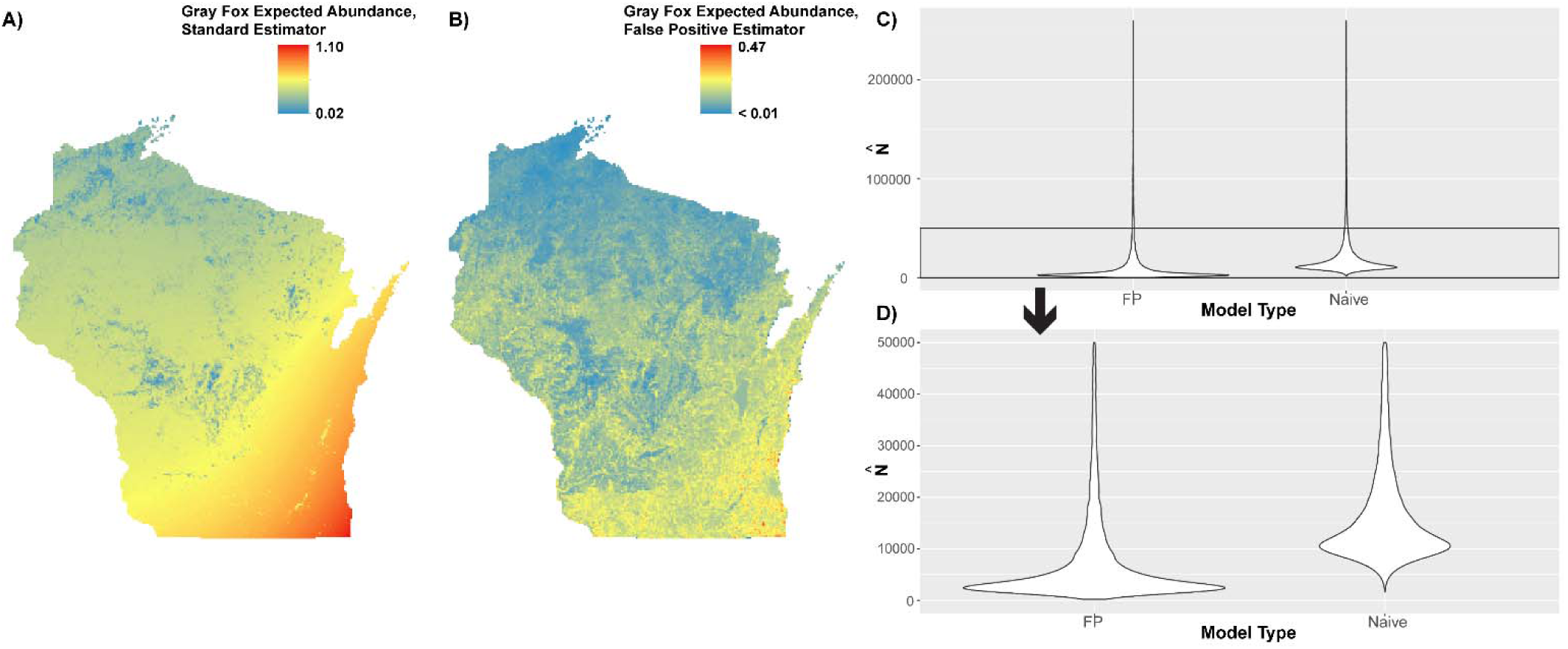
Predictions of gray fox relative abundance across Wisconsin, USA in 2017 using the standard Royle-Nichols estimator (A) and the extended version of the estimator (B) exhibit differences in pattern and granularity; posterior distributions of expected statewide abundance truncated at the upper 95% CI (C) and at 50000 (D) for visualization purposes.

## Discussion

There are several potential reasons that few applied studies to date have fit models to detection-nondetection data that ignore false positive error. Investigators may not typically believe any data is misclassified or may find it difficult to collect additional data to inform model-based differentiation between true and false positive detections. These topics are not the focal area of our study, but the first is probably not often strictly true, and the second is increasingly surmountable. False positive errors have been documented within image, audio, or live observation tasks performed by individual volunteers, machine-learning algorithms, paid professionals, or via crowdsourcing (Miller et al. 2012, Swanson et al. 2016, Abra et al. 2018, Nourazaddeh et al. 2018). Like imperfect detection, false positive error may be the “rule rather than the exception” (Chen et al. 2013) unless all observations have been thoroughly reviewed (Gardiner et al. 2012).

Furthermore, as the number of described protocols for addressing false positive error continues to increase, there are fewer sampling schemes for which false positive error remains inestimable. With respect to the observation confirmation protocol, the requisite data is probably more readily available than ever before. Ecologists are increasingly using automated recording devices for sampling that allow every observation to be reviewed, and technology capable of producing verifiable data has proliferated such that confirmation may often be possible even when recordings are not the primary sampling technique. Any individual with a smartphone has the opportunity to produce a verifiable observation (i.e., a photograph or audio recording) out of an ephemeral observation like a track, sighting, or song.

The more defensible reasons to ignore false positive error are, in our view, primarily practical. These include lack of any described false positive extension for the model of interest, the expectation that false positive error is not going to substantially affect inference or prediction (e.g., because it is sufficiently uncommon), or concerns that a more general model might become too computationally burdensome. We discuss these considerations further below.

The chief benefit of the generalized model description provided here, we hope, is that it makes it easier for investigators to incorporate false positive error within detection-nondetection models of interest. The observation-confirmation structure can be easily applied to essentially any described parametric model for detection-nondetection data by altering the definition of *θ_tp_*. Indeed, its extensibility exceeds our capacity to investigate it here, and we highlight potential avenues for future research. More specific simulation studies are necessary to develop general rules of thumb for optimizing sampling or verification protocols (Clement 2016), particularly for model classes using binary data that were not considered here: presence-background models (Renner et al. 2015), models that integrate occurrence with other data types (Chandler and Clark 2014, Zipkin et al. 2017), or dynamic models: our intuition is that although accounting for false positive error is rarely an incorrect choice, certain models are more sensitive to false positive error than others (see below). Similarly, further evaluation is needed to inform the practical implementation of other protocols for false positive error (again, also extensible: Appendix S1) across different model types. Finally, we note that the general structure for false positive error within binary data—*y_i,j_*~ Bernoulli (*θ_tp_*+ *θ_fp_*) or Bernoulli (*θ_tp_* + *θ_fp_* – [*θ_tp_*×*θ_fp_*])—can also be extended to other data types. For example, a count of organisms *y_i,j_* could be considered distributed as Poisson (λ*_tp_* + λ*_fp_*) assuming that each organism is counted once: developing sampling protocols that enable identification of these parameters is a clear future research need.

Our simulation results demonstrate that rates of misclassification heretofore considered sufficiently accurate (e.g., 5% of classifications are false positive) can induce potentially problematic levels of bias in estimates of abundance or arrival time. However, a key point emerging from our study is that intuiting whether false positive error is likely to substantively influence results or not is tricky. There are two reasons. First, false positive error affects estimation and inference in several varying ways. Consistent with previous work (e.g., Miller et al. 2011), false positive error tended to inflate estimates of species distribution or abundance, and negatively bias estimates of arrival if false positives occur prior to species arrival. One potential justification for ignoring false positive error in the face of potential bias might be that the research objectives can be achieved with ordinal or proportional inference, like determining the direction of a temporal trend identifying a general spatial pattern (Guillera-Arroita et al. 2015, Cruickshank et al. 2019), or that the research objectives prioritize minimizing uncertainty more than bias. However, we also demonstrate that false positive error can sometimes *increase* estimate uncertainty (e.g., arrival time) and can distort predictive patterns or trends (as seen in the application). Neither finding is surprising given that temporally distinct false positives are likely to increase the variance in observed arrival times and that spatially distinct false positives inherently alter patterns in observed occurrence or abundance. What they collectively suggest is that investigators should think more carefully and broadly about how false positive error might impact estimation of state variables or parameters of interest before assuming said error is inconsequential.

Secondly, estimator sensitivity to false positive error is further dependent upon several other factors. As with previous studies (e.g., Ruiz-Gutiérrez et al. 2016), we found that estimator bias tended to increase when there were more false positives within the dataset. However, sensitivity to false positive error is also likely to depend upon the degree of spatial or temporal overlap between true and false positive observations (simulations suggest greater bias with less overlap, and presumably less overlap also distorts inference regarding spatiotemporal patterns more strongly), how efficiently the state process of interest is observed (distribution estimates were more biased when the conditional probability of true detection was lower), and several other attributes of and interactions between the state process, sampling design, and object of inference. For example, the effect of false positives upon occupancy estimates is less pronounced when a species is very widespread (e.g., Clare et al. 2019), and presumably, the effect of false positives upon estimated arrival times depends partially upon when sampling is initiated. There is also likely to be sensitivity to model form and structure. Unlike occupancy models (e.g., Clare et al. 2019), the RN model exhibited consistent bias across a (limited) range of abundances. Moreover, the range of potential bias for the RN model induced by false positives appears to exceed the range of potential bias exhibited by occupancy models: the case study suggests that at relatively high rates of false positive error, the RN model’s bias can be extreme.

Other considerations include the complexity of the fitted functional response and model assumptions pertaining to imperfect detection. Complex responses (e.g., splines) may be more distorted by false positive error than monotonic response shapes (Fernandes et al. 2019). Similarly, we expect that models assuming imperfect detection are particularly sensitive to false positive error because species are both observed in more places, and observation patterns become more heterogeneous and sparse (Kéry and Royle 2016). In contrast, a 5% false positive rate can induce at most a 5% absolute bias in finite-sample estimates in area of occurrence assuming perfect detection, and has no influence on prevalence within a presence-background model because the parameter is inestimable. Given such variability that, for example, a 5% false positive rate could induce either practically no bias or 40% relative bias in estimated population size, it is clear that assuming false positive error is inconsequential actually implies several distinct assumptions. It is also difficult to see how general blanket definitions for data sufficiency (e.g., 95% classification accuracy) can be effective across different model types, species, or sampling protocols without being extremely stringent (e.g., 99% accuracy).

An underappreciated benefit of explicitly modeling false positive error is that it eases the burden associated with thinking about several potential drivers of bias. It is not that false positive models are completely insensitive to the prevalence of error or its patterns or sampling/state considerations. In fact, it is worth emphasizing that accounting for false positive error does not correct any other model or data deficiencies: small sample size situations (e.g., low detection probability, few sites) that are problematic for standard estimators are just as problematic for their false positive extensions. Still, as we have argued previously, we believe that in many cases variability in estimator bias is typically muted once a model accounts for false positives (Clare et al. 2019). Once the low to moderate amounts of false positive error exhibiting constant or univariate patterns in our simulations were accounted for, the number of confirmed samples or the structure of the model for false positive error had limited influence on estimator bias. For species and sampling schemes with relatively limited amounts of false positive error, there may be little practical benefit to confirming more than, say, 50 observations, and a constant model for false positive error may be sufficient (assuming the model for *θ_tp_* is adequately specified) because there is either limited variation to explain or limited benefit to trying to explain more variation.

For species or sampling schemes with a greater prevalence of error (as was the case with the gray fox in our case study), a larger confirmation sample is likely to be necessary (Ruiz-Gutiérrez et al. 2016, Chambert et al. 2018). Furthermore, as the proportion of false positive error increases and there is potentially more variation in false positive error to explain, a larger confirmation sample allows one to fit more complex parameterizations for false positives using splines, zero-inflation terms, or functions based upon latent variables (e.g., the abundance of look-alike species or the species of question). The efficacy of more complex models for false positive error (and the adequacy of simpler models) across a realistic range of sampling conditions is an important question. In part, we used relatively simple models for simulation simply because it is not clear what constitutes a realistic or adequate generating model for false positive error.

Relatedly, the degree to which accounting for false positive error reduces bias also depends upon how well the model for true positives approximates the generating process (McClintock et al. 2010b, Miller et al. 2015). In principal, the model-based solutions for false positives operate by probabilistically distinguishing true and false positives: estimation of each part of the mixture is sensitive to the specification of the other. The case study represents a particularly problematic dataset. The environmental associations of the gray fox were poorly understood and there was a large proportion of false positive error. Even worse, this error was associated with two other generalist species with poorly delineated habitat associations that also, across much of the state, are believed to exist at greater density (e.g., analysis associated with Clare et al. 2016). We almost certainly did not fit the true models for gray fox true and false positive detections, and we do not know which population estimate or predictive surface is closer to the truth. However, we have higher confidence in the estimate accounting for false positive error because its predictions more closely resemble the patterns described by previous survey efforts (Peterson et al. 1977) and because results here and elsewhere (Miller et al. 2015) suggest that there is greater marginal benefit associated with relaxing the assumption of no false positive error than marginal loss associated with model misspecification.

Accounting for false-positive error adds more parameters and added dimensions to the objective function of interest—it is somewhat slower to fit extended models, and there is an additional component that might be subject to model selection practices. However, we think there are several ways to alleviate any additional computational burden. First, while we describe the extensions using a complete data likelihood because it makes extensibility more intuitive, neither the complete data likelihood nor Bayesian techniques are necessary for implementation. It is also possible to integrate distinct likelihoods for ***y*** and ***v*** over possible values for any latent variables to construct faster estimators using maximum likelihood or Hamiltonian-MCMC (see example with the RN model in Appendix S4). Secondly, because the generating process for *θ_fp_* is the same regardless of how the model for *θ_tp_* is specified, we think it is reasonable to try to address any model selection for false positive error within a simpler model structure for *θ_tp_*: for example, selecting the model structure for false positives within a standard occupancy estimator. Finally, defining a prior distribution for *θ_fp_* informed by the estimates of a simpler model may make estimation faster (if slightly less efficient) than incorporating confirmed data directly into the likelihood. This is likely to produce larger computational gains when the number of confirmed samples is sizable. There are, however, a few caveats. First, the confirmation data contains information useful for inferring specific latent states, and this information is lost when using a prior distribution for *θ_fp_* (as it is when using a calibration protocol). Secondly, estimation of *θ_fp_* is likely sensitive to the specification of *θ_tp_*: arrival and RN models are structurally very similar to occupancy models, but transferability between occupancy models and very distinct model types is less certain. Finally, it is unclear how transferable *θ_fp_* is likely to be if it is modeled as varying in more complex ways than considered here.

In whole, we believe we have demonstrated that it is possible and tractable to address false positive error across a broader range of estimation problems than previously considered. We conclude by noting that the effects of false positive error, the strategies for dealing with it, and the limitations of these strategies largely mirror considerations associated with imperfect detection. The importance of imperfect detection, the practicality of collecting the extra data needed to estimate it within models relative to trying to control for it using design-based solutions, and the stability of models accounting for imperfect detection have all been debated, and these considerations are all similarly germane to false positive error. It is now generally accepted that imperfect detection needs to be accounted for to achieve certain objectives and that statistical techniques to do so are important ecological tools (Guillera-Arroita et al. 2015). It has been argued that false positive error deserves similar wide consideration (Miller et al. 2015). The generalizable model structure described here, we hope, will accelerate the adoption of techniques to limit its effects while providing investigators the flexibility to fit the models they are interested in fitting.

## Acknowledgments

Support for this research was provided by NASA ESSF NNX16AO61H and NASA Ecological Forecasting grant NNX14AC36G. We thank the associate editor and 2 anonymous reviewers for constructive comments on previous drafts.

## Appendix S1. Other model extensions

### Extension of other false-positive models

Despite our focus on leveraging observation-confirmation to deal with false positive error, it is important to note that other strategies for estimating false positive parameters can also be effective across different model types. We describe here alternative formulations following different described protocols for dealing with false positive error when estimating relative abundance using the model of Royle and Nichols (2003). The base model can be expressed hierarchically as:

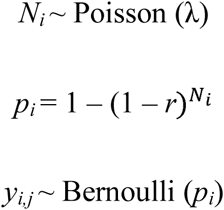

The description for the observation-confirmation extension described in the main text is:

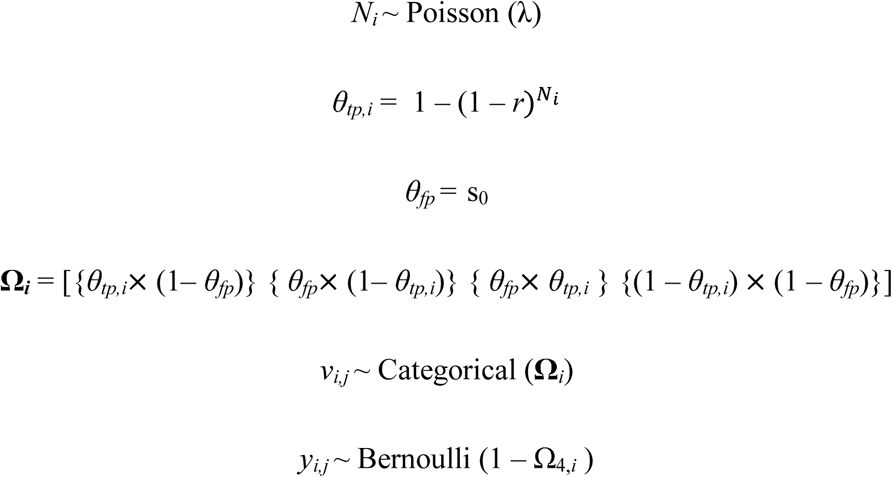

As noted in the main text, a distinctive feature of the observation-confirmation protocol is that true and false positives are not necessarily mutually exclusive in space or space/time. The alternative protocols do require the assumption that true and false positive observations are distinct in space (i.e., false positive observations can only occur in places where true positive observations are impossible. This may not reflect the reality of the sampling itself (i.e., there may well be true and false positive observations at the same location or the same location at the same time), but is a constraint imposed by the nature of the information that these protocols make available. ***As noted in the discussion, we do not know how effectively the other protocols extend to other model structures…this is an open research question that deserves attention*.**

For the RN model, perhaps the most logical way to make *θ_fp_* and *θ_tp_* mutually exclusive is to constrain *θ_fp_* to only occur at locations where abundance is zero and true positives cannot occur. A formulation in the spirit of Royle and Link (2006) follows:

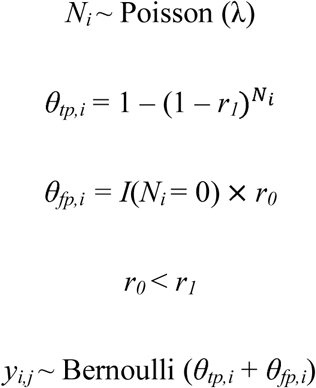

Here, *I*(*N_i_* = 0) denotes an indicator function for whether the abundance of site *i* = 0, and *r_0_* denotes the probability of falsely detecting an organism at a location with zero abundance. The constraint *r_0_* < *r_1,_* where *r_1_* is the probability of detecting a single present organism, mirrors the constraint in the original model that *p_10_* < *p_11_*.

Within the site-confirmation protocol (e.g., Chambert et al. 2015), the observed observations *y_i,j_* follow a categorical distribution pertaining to whether no species was observed, a species was observed but not confirmed, and a species was unambiguously observed. The parameter *b* describes the conditional probability that a positive observation is unambiguous. A Royle-Nichols variant of the site-confirmation protocol can be described as:

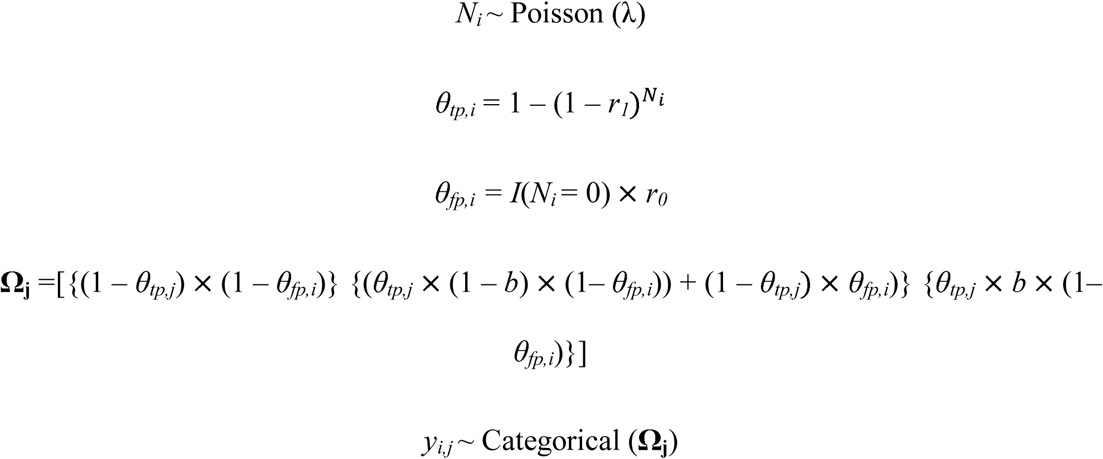

One variant of the site-confirmation protocol, the multiple detection methods model (Miller et al. 2011, Chambert et al. 2015), assumes that one sampling method (M1) generates ambiguous positive detections *y_i,j_* and a second independent method (M2) operating during different occasions *s* always provides unambiguous positive detections *w_i,s_*. The distinct probabilities of truly detecting a single animal for M1 and M2 can respectively be described as *r_1_* and *r_2_*. The hierarchical formulation is then:

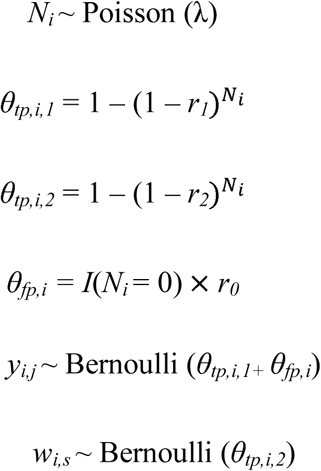

Finally, following the calibration protocol (Chambert et al. 2015), an investigator might have collected reference detection data under experimental conditions in which the state is known (or at least, the exclusive conditions for *θ_tp_* and *θ_fp_* are known): *x_1_* and *x_0_* respectively denote the total number of true and false positive detections in the reference data set, and *n_1_* and *n_0_* the number of trials in which true positive detections were possible or not—an example trial might be the presentation of a single image or recording in which the species of interest is present or not present. One way to specify the Royle-Nichols model hierarchically following the calibration protocol is:

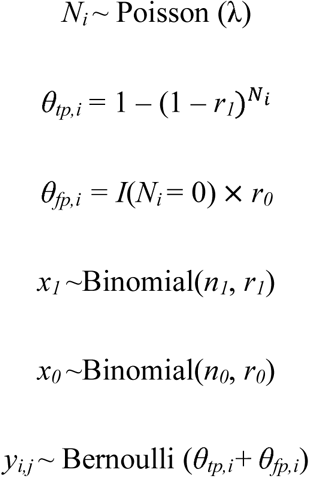

The key to implementing these different protocols within different model structures is only slightly more complex than extending the observation-confirmation protocol: one must redefine *θ_tp,i_* following the original model description, and redefine *θ_fp,i_* so that is exclusive of *θ_tp,i_*, and alter the statements associated with the auxiliary data appropriately. For example, to define extensions for the arrival model following the protocols above instead of the RN model, define *θ_tp,i,j_* = *z_i_* × *p_1_* × I(*j > x_i_*), *θ_fp,i,,j_ =* (1 – (*z_i_* × I(*j > x_i_*)) × *p_0_*, and throughout, replace *r_0_*, *r_1_*, and *r_2_* with, respectively, (say) *p_0_*, *p_1_*, and *p_2_*.

### Observation Confirmation Protocol for the Spatial Royle-Nichols Model

The spatial Royle-Nichols model (Ramsay et al. 2015) uses *z_i_* to denote whether individuals *i* = 1,2…M exist within a geographic space ||*S*|| with probability *ψ*. The state variable of interest, population size *N* in ||*S*||, is estimated as 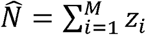, and population density is derived as 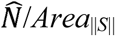. Individuals have distinct activity centers located within ||*S*|| and the coordinates of these activity centers are denoted as *s_i_*; individuals are detected at any of *j* detectors on given sampling occasions *k* with probability *p_i,j_*. The unconditional probability of detection is a function of whether an individual exists, the distance between an individual’s latent activity center and the location of the detector, *d_i,j_*, and the parameters *g_0_* and σ that respectively relate to the probability of individual detection at *d_i,j_*= 0 and the rate at which individual encounter probability decays, and can be expressed as 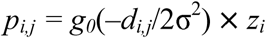. Individuals are not distinguished, so these parameters are inferred by marginalizing across the latent individual encounter histories at a specific detector such that the unconditional probability of detection is described as 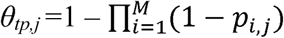. The hierarchical likelihood is:

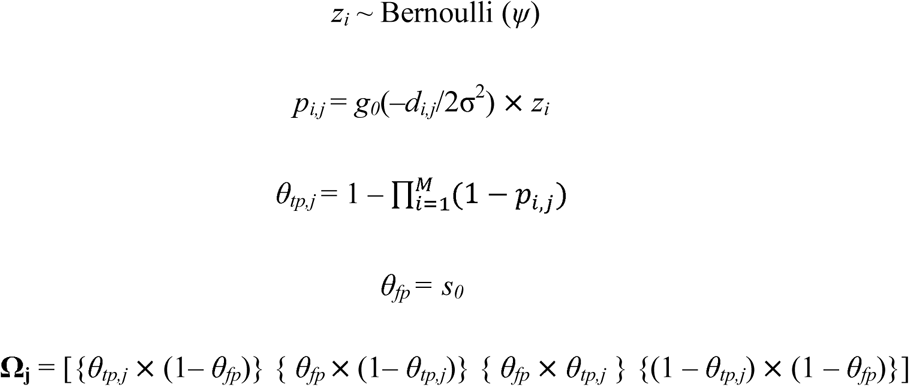

Here, *v_j,k_* ~Categorical (Ω**_j_**) and *y_j,k_* ~ Bernoulli (1 – Ω_4,j_). Critically, *θ_tp,j_* has no support at 0, and as a result, without re-specifying the SRN model or one of the false positive protocols, site-confirmation or calibration approaches to false positive error are impossible to implement. Although not directly considered within the manuscript, code required to simulate and fit extended SRN models is found in within Appendix S4. As a general proof of concept that the model is sensitive to false positive error and an extended version can reduce bias, we present very limited simulation results here.

We simulated 100 replicate datasets to demonstrate proof of concept. Sampling parameters included a population size of 50 organisms; ||*S*|| defined as a 20 × 20 unit square; detection parameters *g_0_* = 0.15 and σ = 0.5; 196 detector locations within a 14 × 14 square grid with 1 unit spacing, and 20 sampling intervals: only the location of individual activity centers varied across simulation replicates. False positive observations (as 10% of all detections) and a verification sample were simulated following practices described in the main text; the size of the verification sample varied as{10 20 30 40 50} replicated 20 times each. We compared the standard model and the false positive extension on the basis of relative bias of 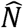. The standard estimator was strongly biased (relative bias = 0.81) at a 10% false positive rate; the extended model was not unbiased—perhaps due to small sample size considerations—but exhibited better performance (relative bias = 0.25). Clearly more research is needed to understand the general sensitivity of the SRN model to false positive error. Given the narrow range of parameter space within which the base model is unbiased (Ramsey et al. 2015), we expect that it is more likely to see usage as a component within integrated models (Sun et al. 2019), and we suggest that it may be more fruitful to explore sensitivity to false positives within this class of model.

## Appendix S2. Additional details associated w/ model performance

**Table S1.** Scenarios considered when evaluating the Royle-Nichols and extended models. β describes abundance terms, α describes detection terms. ***X_1_*** is a vector of simulated covariates that influences abundance, and for some scenarios, the likelihood of false positive observation.

| Scenario | $\beta_0$ | $\beta_1$ | $\alpha_0$ | $\alpha_1$ | False Positive Model |
| --- | --- | --- | --- | --- | --- |
| 1 | 0 | 1 | -1.73 | 1 | $\theta_{fp} = 0.10$ of $\theta_{fp} + \theta_{tp}$ |
| 2 | 0 | 1 | -1.73 | 1 | $\theta_{fp} = 0.05$ of $\theta_{fp} + \theta_{tp}$ |
| 3 | 0 | 1 | -1.73 | 1 | $\theta_{fp} = 0.01$ of $\theta_{fp} + \theta_{tp}$ |
| 4 | -1.50 | 1 | -1.73 | 1 | $\theta_{fp} = 0.10$ of $\theta_{fp} + \theta_{tp}$ |
| 5 | -1.50 | 1 | -1.73 | 1 | $\theta_{fp} = 0.05$ of $\theta_{fp} + \theta_{tp}$ |
| 6 | -1.50 | 1 | -1.73 | 1 | $\theta_{fp} = 0.01$ of $\theta_{fp} + \theta_{tp}$ |
| 7 | -1.50 | 1 | -1.73 | 1 | $\text{logit}(\theta_{fp,i}) = -6 + 1X_{1,i}$ |
| 8 | -1.50 | 1 | -1.73 | 1 | $\text{logit}(\theta_{fp,i}) = -6 - 1X_{1,i}$ |

**Table S2.** Mean error for parameters and relative bias of finite-sample population size for the standard and extended RN models across the scenarios considered.

| Estimator | Scenario | Mean Error (Absolute Bias) |  |  |  | Relative Bias |
| --- | --- | --- | --- | --- | --- | --- |
| | | $\hat{\beta}_0$ | $\hat{\beta}_1$ | $\hat{\alpha}_0$ | $\hat{\alpha}_1$ | $\hat{N}^*$ |
| Standard | 1 | 0.71 | -0.13 | -0.58 | -0.2 | 0.86 |
|  | 2 | 0.42 | -0.08 | -0.35 | -0.12 | 0.45 |
|  | 3 | 0.1 | -0.02 | -0.08 | -0.03 | 0.1 |
|  | 4 | 0.71 | -0.2 | -0.48 | -0.19 | 0.79 |
|  | 5 | 0.37 | -0.1 | -0.28 | -0.12 | 0.38 |
|  | 6 | 0.08 | -0.03 | -0.07 | -0.01 | 0.09 |
|  | 7 | 0.28 | 0 | -0.23 | -0.1 | 0.37 |
|  | 8 | 0.61 | -0.32 | -0.35 | -0.16 | 0.51 |
| False Positive Extension | 1 | -0.01 | 0.01 | -0.01 | 0 | 0.02 |
|  | 2 | -0.02 | 0.01 | 0 | 0 | 0.01 |
|  | 3 | -0.01 | -0.01 | 0 | 0 | 0.01 |
|  | 4 | -0.02 | -0.01 | -0.01 | 0.01 | 0.02 |
|  | 5 | -0.06 | 0.02 | 0 | 0 | 0.01 |
|  | 6 | -0.03 | 0.01 | -0.01 | 0.02 | 0.01 |
|  | 7 | -0.03 | 0 | -0.01 | 0.01 | 0.02 |
|  | 8 | -0.06 | 0 | 0 | 0.01 | -0.01 |
|  | 1<sup>A</sup> | -0.04 | 0.02 | 0 | 0.01 | 0 |
|  | 2<sup>A</sup> | -0.04 | 0.02 | 0.01 | 0.01 | 0 |
|  | 3<sup>A</sup> | -0.02 | 0 | 0.01 | 0 | 0 |
|  | 4<sup>A</sup> | -0.02 | -0.01 | -0.01 | 0.01 | 0.02 |
|  | 5<sup>A</sup> | -0.06 | 0.02 | 0 | 0 | 0.01 |
|  | 6<sup>A</sup> | -0.03 | 0.01 | 0 | 0.02 | 0.01 |
|  | 7<sup>A</sup> | -0.03 | -0.01 | -0.01 | 0.01 | 0.01 |
|  | 8<sup>A</sup> | -0.07 | 0 | 0 | 0.01 | -0.01 |
|  | 7<sup>B</sup> | -0.01 | 0.04 | -0.02 | 0 | 0.06 |
|  | 8<sup>B</sup> | -0.06 | -0.03 | 0.01 | 0.01 | -0.02 |

<sup>A</sup>Fit using an informed prior for $\theta_{fp}$ derived from estimates from an occupancy model following the observation confirmation protocol.
<sup>B</sup>A slightly mis-specified model that assumes $\theta_{fp}$ is constant.

**Table S3.** Frequentist coverage of 95% CRI associated with parameters and finite-sample population size for the standard and extended RN models across the scenarios considered.

| Estimator | Scenario | Coverage |  |  |  |  |
| --- | --- | --- | --- | --- | --- | --- |
| | | $\hat{\beta}_0$ | $\hat{\beta}_1$ | $\hat{\alpha}_0$ | $\hat{\alpha}_1$ | $\hat{N}^*$ |
| Standard | 1 | 0 | 0.403 | 0 | 0.216 | 0 |
|  | 2 | 0.093 | 0.756 | 0.176 | 0.556 | 0.083 |
|  | 3 | 0.84 | 0.913 | 0.87 | 0.89 | 0.866 |
|  | 4 | 0.066 | 0.556 | 0.086 | 0.523 | 0.013 |
|  | 5 | 0.456 | 0.816 | 0.46 | 0.736 | 0.193 |
|  | 6 | 0.906 | 0.926 | 0.9 | 0.876 | 0.9 |
|  | 7 | 0.636 | 0.936 | 0.566 | 0.783 | 0.246 |
|  | 8 | 0.066 | 0.286 | 0.243 | 0.626 | 0.063 |
| False<br>Positive<br>Extension | 1 | 0.983 | 0.95 | 0.95 | 0.93 | 0.95 |
|  | 2 | 0.99 | 0.94 | 0.96 | 0.95 | 0.95 |
|  | 3 | 0.98 | 0.95 | 0.97 | 0.96 | 0.96 |
|  | 4 | 0.97 | 0.94 | 0.96 | 0.95 | 0.96 |
|  | 5 | 0.97 | 0.96 | 0.96 | 0.92 | 0.96 |
|  | 6 | 0.98 | 0.93 | 0.93 | 0.92 | 0.97 |
|  | 7 | 0.97 | 0.93 | 0.94 | 0.953 | 0.95 |
|  | 8 | 0.98 | 0.92 | 0.96 | 0.96 | 0.92 |
|  | 1<sup>A</sup> | 0.98 | 0.95 | 0.95 | 0.94 | 0.92 |
|  | 2<sup>A</sup> | 0.99 | 0.95 | 0.96 | 0.95 | 0.96 |
|  | 3<sup>A</sup> | 0.99 | 0.94 | 0.96 | 0.96 | 0.96 |
|  | 4<sup>A</sup> | 0.98 | 0.94 | 0.97 | 0.95 | 0.96 |
|  | 5<sup>A</sup> | 0.98 | 0.96 | 0.96 | 0.93 | 0.94 |
|  | 6<sup>A</sup> | 0.98 | 0.92 | 0.92 | 0.92 | 0.98 |
|  | 7<sup>A</sup> | 0.973 | 0.94 | 0.94 | 0.96 | 0.95 |
|  | 8<sup>A</sup> | 0.973 | 0.93 | 0.95 | 0.95 | 0.92 |
|  | 7<sup>B</sup> | 0.97 | 0.93 | 0.94 | 0.96 | 0.9 |
|  | 8<sup>B</sup> | 0.97 | 0.91 | 0.96 | 0.953 | 0.92 |

<sup>A</sup>Fit using an informed prior for $\theta_{fp}$ derived from estimates from an occupancy model following the observation confirmation protocol.
<sup>B</sup>A slightly mis-specified model that assumes $\theta_{fp}$ is constant.

**Table S4.** Root mean squared error for parameters and relative bias of finite-sample population size for the standard and extended RN models across the scenarios considered.

|  |  | RMSE |  |  |  |  |
| --- | --- | --- | --- | --- | --- | --- |
| Estimator | Scenario | $\hat{\beta}_0$ | $\hat{\beta}_1$ | $\hat{\alpha}_0$ | $\hat{\alpha}_1$ | $\widehat{N}^*$ |
| Standard | 1 | 0.71 | 0.14 | 0.58 | 0.2 | 280 |
|  | 2 | 0.42 | 0.09 | 0.36 | 0.13 | 150 |
|  | 3 | 0.1 | 0.07 | 0.11 | 0.07 | 39 |
|  | 4 | 0.786 | 0.21 | 0.49 | 0.21 | 59 |
|  | 5 | 0.377 | 0.14 | 0.28 | 0.16 | 28 |
|  | 6 | 0.173 | 0.12 | 0.13 | 0.12 | 9 |
|  | 7 | 0.29 | 0.11 | 0.24 | 0.15 | 27 |
|  | 8 | 0.61 | 0.32 | 0.36 | 0.2 | 37 |
| False<br>Positive<br>Extension | 1 | 0.11 | 0.07 | 0.1 | 0.07 | 33 |
|  | 2 | 0.11 | 0.06 | 0.1 | 0.07 | 29 |
|  | 3 | 0.1 | 0.06 | 0.08 | 0.06 | 25 |
|  | 4 | 0.17 | 0.12 | 0.12 | 0.11 | 7.2 |
|  | 5 | 0.17 | 0.12 | 0.12 | 0.11 | 6.65 |
|  | 6 | 0.16 | 0.12 | 0.12 | 0.11 | 6 |
|  | 7 | 0.17 | 0.13 | 0.12 | 0.11 | 7.2 |
|  | 8 | 0.19 | 0.13 | 0.12 | 0.11 | 6.64 |
|  | 1<sup>A</sup> | 0.12 | 0.07 | 0.1 | 0.07 | 32 |
|  | 2<sup>A</sup> | 0.11 | 0.06 | 0.1 | 0.07 | 28 |
|  | 3<sup>A</sup> | 0.1 | 0.06 | 0.09 | 0.06 | 25 |
|  | 4<sup>A</sup> | 0.18 | 0.12 | 0.12 | 0.11 | 7.37 |
|  | 5<sup>A</sup> | 0.17 | 0.12 | 0.12 | 0.11 | 6.92 |
|  | 6<sup>A</sup> | 0.16 | 0.12 | 0.12 | 0.11 | 6.45 |
|  | 7<sup>A</sup> | 0.17 | 0.13 | 0.13 | 0.11 | 7.33 |
|  | 8<sup>A</sup> | 0.19 | 0.13 | 0.12 | 0.11 | 6.89 |
|  | 7<sup>B</sup> | 0.17 | 0.13 | 0.12 | 0.1 | 8.12 |
|  | 8<sup>B</sup> | 0.19 | 0.14 | 0.12 | 0.11 | 6.85 |

<sup>A</sup>Fit using an informed prior for $\theta_{fp}$ derived from estimates from an occupancy model following the observation confirmation protocol.
<sup>B</sup>A slightly mis-specified model that assumes $\theta_{fp}$ is constant.

**Table S5.** Scenarios considered when evaluating the arrival model of Roth et al. (2014) and extended models. *β* describes occupancy terms, *α* describes detection terms. ***X_1_*** is a vector of simulated covariates that influences abundance, and for some scenarios, the likelihood of false positive observation. *φ* denotes the simulated occasion of arrival. ***X_1_*** is a vector of simulated covariates that influences the probability of occupancy, and for some scenarios, the likelihood of false positive observation.

| Scenario | $\beta_0$ | $\beta_1$ | $\alpha_0$ | $\alpha_1$ | $\phi$ | Start of False Positives | False Positive Model |
| --- | --- | --- | --- | --- | --- | --- | --- |
| 1 | 0 | 0.5 | -2 | 0.5 | 6 | >0 | $\theta_{fp} = 0.10$ of $\theta_{fp} + \theta_{tp}$ |
| 2 | 0 | 0.5 | -2 | 0.5 | 6 | >0 | $\theta_{fp} = 0.05$ of $\theta_{fp} + \theta_{tp}$ |
| 3 | 0 | 0.5 | -2 | 0.5 | 6 | >0 | $\theta_{fp} = 0.01$ of $\theta_{fp} + \theta_{tp}$ |
| 4 | 0 | 0.5 | -2 | 0.5 | 6 | 5 | $\text{logit}(\theta_{fp,i}) = -6 + 1X_{1,i}$ |
| 5 | 0 | 0.5 | -2 | 0.5 | 6 | 5 | $\text{logit}(\theta_{fp,i}) = -6 - 1X_{1,i}$ |
| 6 | 0 | 0.5 | -2 | 0.5 | 6 | 3 | $\text{logit}(\theta_{fp,i}) = -6 + 1X_{1,i}$ |
| 7 | 0 | 0.5 | -2 | 0.5 | 6 | 3 | $\text{logit}(\theta_{fp,i}) = -6 - 1X_{1,i}$ |
| 8 | 0 | 0.5 | -1 | 0.5 | 6 | 5 | $\text{logit}(\theta_{fp,i}) = -6 + 1X_{1,i}$ |
| 9 | 0 | 0.5 | -1 | 0.5 | 6 | 5 | $\text{logit}(\theta_{fp,i}) = -6 - 1X_{1,i}$ |

**Table S6.**
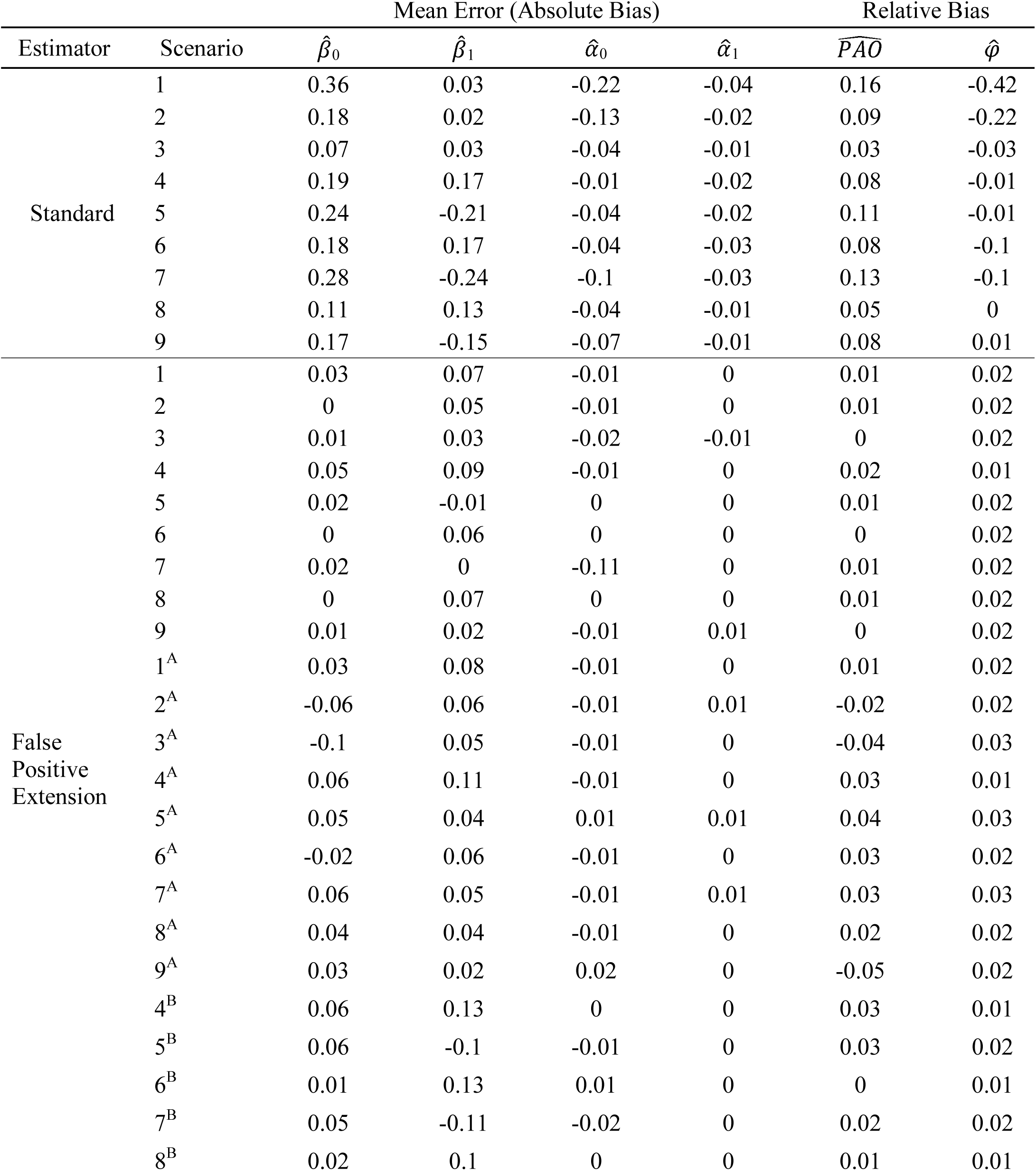
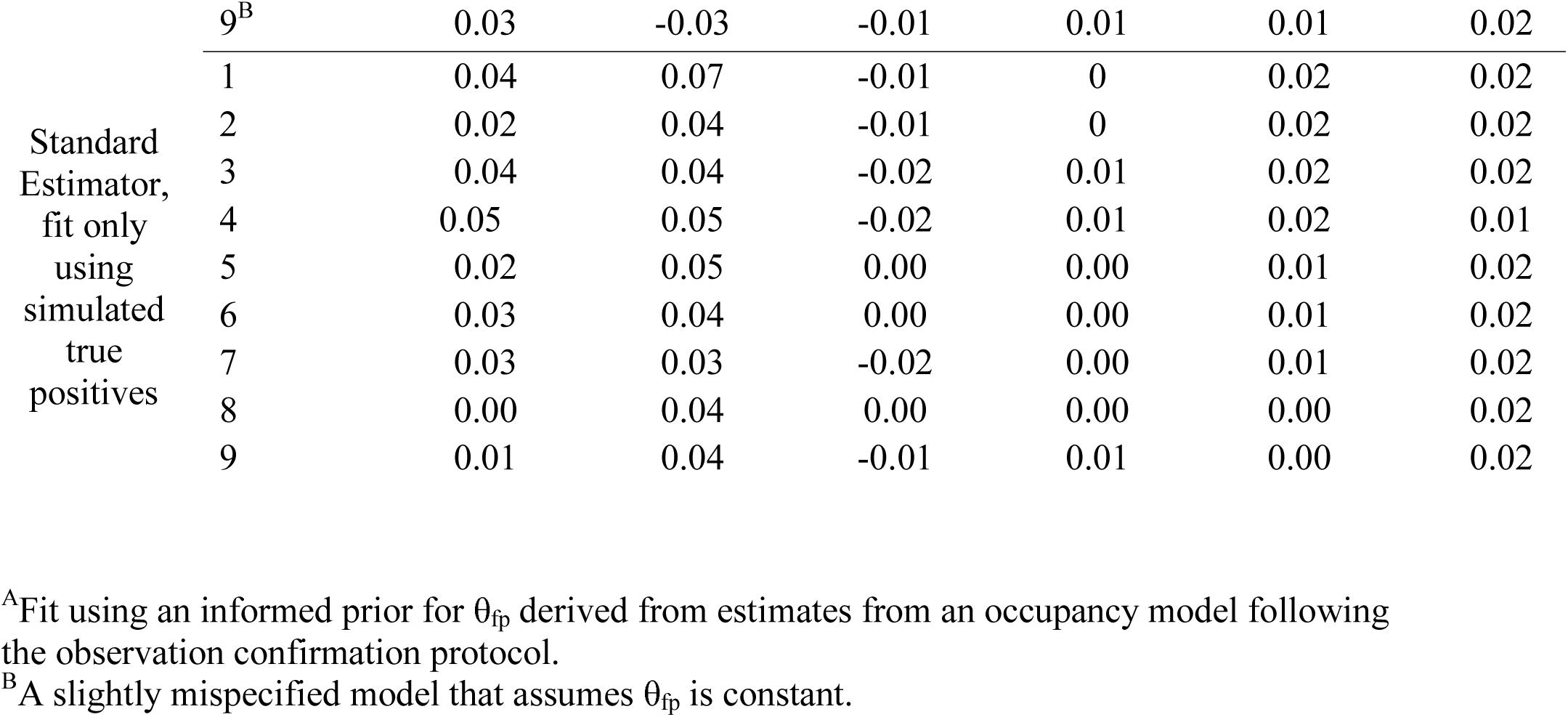
Estimator error and relative bias associated with parameters using the standard and extended arrival occupancy models across the scenarios in table S5.

**Table S7.** Frequentist coverage of 95% CRI associated with parameters using the standard and extended arrival occupancy models across the scenarios in table S5.

| Estimator | Scenario | Coverage |  |  |  |  |  |
| --- | --- | --- | --- | --- | --- | --- | --- |
| | | $\hat{\beta}_0$ | $\hat{\beta}_1$ | $\hat{\alpha}_0$ | $\hat{\alpha}_1$ | $\widehat{PAO}$ | $\hat{\phi}$ |
| Standard | 1 | 0.54 | 0.94 | 0.41 | 0.9 | 0.25 | 0.1 |
|  | 2 | 0.84 | 0.94 | 0.71 | 0.93 | 0.64 | 0.35 |
|  | 3 | 0.94 | 0.93 | 0.94 | 0.95 | 0.93 | 0.82 |
|  | 4 | 0.85 | 0.9 | 0.92 | 0.92 | 0.65 | 0.91 |
|  | 5 | 0.81 | 0.76 | 0.91 | 0.94 | 0.53 | 0.93 |
|  | 6 | 0.84 | 0.87 | 0.94 | 0.93 | 0.71 | 0.5 |
|  | 7 | 0.67 | 0.74 | 0.85 | 0.97 | 0.4 | 0.53 |
|  | 8 | 0.89 | 0.86 | 0.87 | 0.91 | 0.26 | 0.92 |
|  | 9 | 0.82 | 0.84 | 0.78 | 0.9 | 0.08 | 0.95 |
| False<br>Positive<br>Extension | 1 | 0.96 | 0.93 | 0.95 | 0.96 | 0.97 | 0.93 |
|  | 2 | 0.96 | 0.93 | 0.96 | 0.95 | 0.95 | 0.95 |
|  | 3 | 0.95 | 0.92 | 0.97 | 0.95 | 0.96 | 0.94 |
|  | 4 | 0.95 | 0.94 | 0.92 | 0.96 | 0.94 | 0.93 |
|  | 5 | 0.96 | 0.95 | 0.93 | 0.95 | 0.94 | 0.93 |
|  | 6 | 0.95 | 0.93 | 0.95 | 0.94 | 0.97 | 0.96 |
|  | 7 | 0.94 | 0.94 | 0.95 | 0.97 | 0.96 | 0.94 |
|  | 8 | 0.95 | 0.93 | 0.95 | 0.95 | 0.97 | 0.90 |
|  | 9 | 0.96 | 0.95 | 0.93 | 0.96 | 0.97 | 0.94 |
|  | 1<sup>A</sup> | 0.97 | 0.93 | 0.97 | 0.95 | 0.97 | 0.94 |
|  | 2<sup>A</sup> | 0.95 | 0.93 | 0.96 | 0.94 | 0.94 | 0.94 |
|  | 3<sup>A</sup> | 0.91 | 0.93 | 0.97 | 0.96 | 0.91 | 0.92 |
|  | 4<sup>A</sup> | 0.94 | 0.94 | 0.93 | 0.95 | 0.96 | 0.94 |
|  | 5<sup>A</sup> | 0.92 | 0.94 | 0.95 | 0.94 | 0.96 | 0.92 |
|  | 6<sup>A</sup> | 0.93 | 0.95 | 0.96 | 0.93 | 0.95 | 0.95 |
|  | 7<sup>A</sup> | 0.94 | 0.94 | 0.95 | 0.94 | 0.96 | 0.94 |
|  | 8<sup>A</sup> | 0.92 | 0.94 | 0.94 | 0.96 | 0.96 | 0.90 |
|  | 9<sup>A</sup> | 0.94 | 0.93 | 0.94 | 0.95 | 0.96 | 0.93 |
|  | 4<sup>B</sup> | 0.94 | 0.9 | 0.92 | 0.95 | 0.93 | 0.92 |
|  | 5<sup>B</sup> | 0.94 | 0.9 | 0.93 | 0.95 | 0.92 | 0.93 |
|  | 6<sup>B</sup> | 0.95 | 0.92 | 0.94 | 0.94 | 0.98 | 0.95 |
|  | 7<sup>B</sup> | 0.93 | 0.9 | 0.95 | 0.97 | 0.93 | 0.93 |
|  | 8<sup>B</sup> | 0.93 | 0.91 | 0.95 | 0.95 | 0.93 | 0.91 |
|  | 9<sup>B</sup> | 0.96 | 0.94 | 0.93 | 0.95 | 0.97 | 0.92 |

<sup>A</sup>Fit using an informed prior for $\theta_{fp}$ derived from estimates from an occupancy model following the observation confirmation protocol.
<sup>B</sup>A slightly mis-specified model that assumes $\theta_{fp}$ is constant.

**Table S8.** Root mean squared error associated with parameters using the standard and extended arrival occupancy models across the scenarios in table S5.

| Estimator | Scenario | RMSE |  |  |  |  |  |
| --- | --- | --- | --- | --- | --- | --- | --- |
| | | $\hat{\beta}_0$ | $\hat{\beta}_1$ | $\hat{\alpha}_0$ | $\hat{\alpha}_1$ | $\widehat{PAO}$ | $\hat{\phi}$ |
| Standard | 1 | 0.36 | 0.18 | 0.224 | 0.08 | 0.08 | 25.2 |
|  | 2 | 0.23 | 0.17 | 0.15 | 0.08 | 0.05 | 13.6 |
|  | 3 | 0.17 | 0.16 | 0.1 | 0.08 | 0.03 | 4 |
|  | 4 | 0.23 | 0.21 | 0.09 | 0.08 | 0.04 | 2.3 |
|  | 5 | 0.25 | 0.24 | 0.1 | 0.07 | 0.06 | 2.4 |
|  | 6 | 0.22 | 0.22 | 0.09 | 0.08 | 0.04 | 6.4 |
|  | 7 | 0.3 | 0.26 | 0.12 | 0.08 | 0.07 | 6.5 |
|  | 8 | 0.15 | 0.18 | 0.06 | 0.06 | 0.03 | 1.5 |
|  | 9 | 0.19 | 0.18 | 0.09 | 0.06 | 0.04 | 1.4 |
| False<br>Positive<br>Extension | 1 | 0.16 | 0.19 | 0.09 | 0.08 | 0.03 | 2.76 |
|  | 2 | 0.16 | 0.17 | 0.09 | 0.08 | 0.03 | 2.45 |
|  | 3 | 0.16 | 0.17 | 0.08 | 0.08 | 0.02 | 2.4 |
|  | 4 | 0.18 | 0.18 | 0.1 | 0.08 | 0.03 | 2.45 |
|  | 5 | 0.16 | 0.17 | 0.1 | 0.07 | 0.03 | 2.7 |
|  | 6 | 0.17 | 0.18 | 0.09 | 0.08 | 0.02 | 2.3 |
|  | 7 | 0.17 | 0.17 | 0.09 | 0.08 | 0.03 | 2.6 |
|  | 8 | 0.13 | 0.15 | 0.06 | 0.06 | 0.01 | 1.54 |
|  | 9 | 0.12 | 0.13 | 0.06 | 0.06 | 0.01 | 1.66 |
|  | 1<sup>A</sup> | 0.17 | 0.2 | 0.09 | 0.08 | 0.03 | 2.84 |
|  | 2<sup>A</sup> | 0.18 | 0.18 | 0.09 | 0.08 | 0.03 | 2.67 |
|  | 3<sup>A</sup> | 0.19 | 0.17 | 0.09 | 0.08 | 0.03 | 2.73 |
|  | 4<sup>A</sup> | 0.19 | 0.19 | 0.10 | 0.08 | 0.03 | 2.45 |
|  | 5<sup>A</sup> | 0.18 | 0.18 | 0.10 | 0.07 | 0.03 | 2.74 |
|  | 6<sup>A</sup> | 0.19 | 0.18 | 0.09 | 0.08 | 0.02 | 2.4 |
|  | 7<sup>A</sup> | 0.19 | 0.18 | 0.09 | 0.08 | 0.03 | 2.7 |
|  | 8<sup>A</sup> | 0.14 | 0.15 | 0.07 | 0.06 | 0.01 | 1.7 |
|  | 9<sup>A</sup> | 0.14 | 0.14 | 0.06 | 0.06 | 0.01 | 1.8 |
|  | 4<sup>B</sup> | 0.18 | 0.2 | 0.1 | 0.07 | 0.03 | 2.4 |
|  | 5<sup>B</sup> | 0.16 | 0.19 | 0.1 | 0.07 | 0.03 | 2.7 |
|  | 6<sup>B</sup> | 0.18 | 0.2 | 0.09 | 0.08 | 0.02 | 2.3 |
|  | 7<sup>B</sup> | 0.17 | 0.19 | 0.09 | 0.08 | 0.03 | 2.6 |
|  | 8<sup>B</sup> | 0.13 | 0.16 | 0.06 | 0.06 | 0.01 | 1.5 |
|  | 9<sup>B</sup> | 0.12 | 0.14 | 0.06 | 0.06 | 0.01 | 1.7 |

<sup>A</sup>Fit using an informed prior for $\theta_{fp}$ derived from estimates from an occupancy model following the observation confirmation protocol.
<sup>B</sup>A slightly misspecified model that assumes $\theta_{fp}$ is constant.

**Figure S1.**
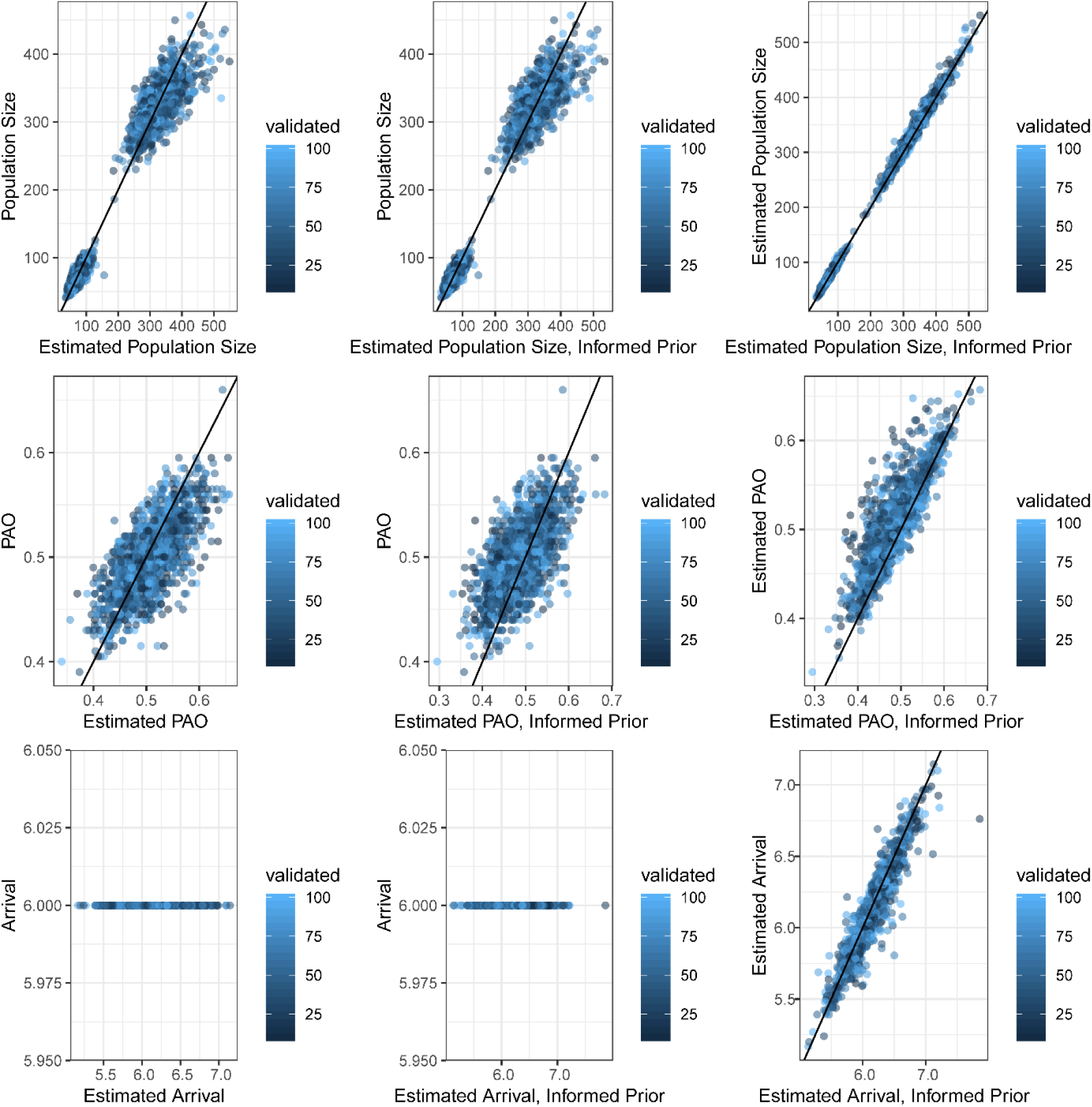
Correlation between true values, estimates made when incorporating confirmed observations, and estimates made when using an informed prior rather than incorporating confirmed observations in the likelihood.

**Figure S2.**
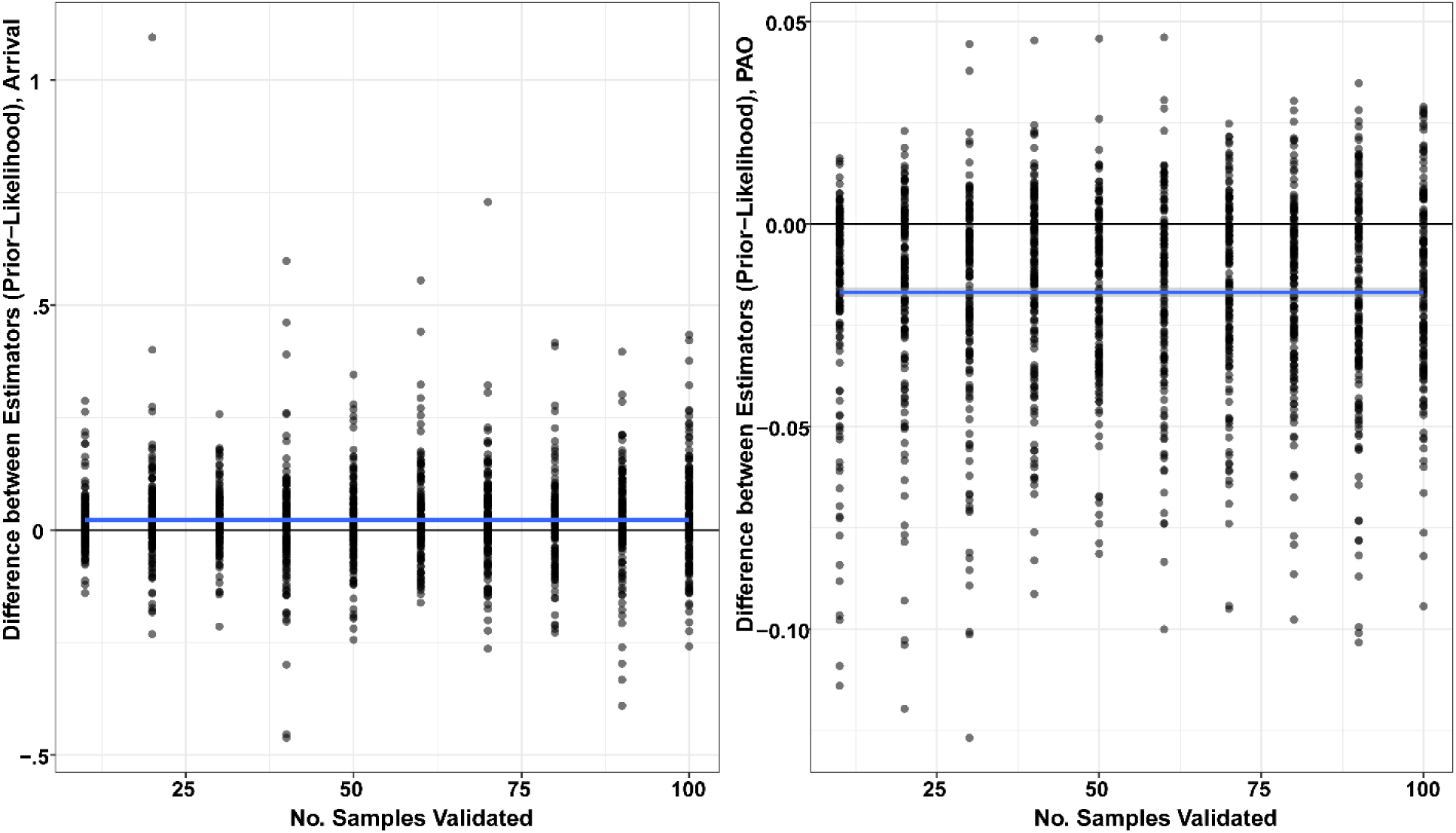
Noise in the correlations between estimates of PAO and arrival generated when including confirmed data in the likelihood vs. using an informed prior is not related to the size of the confirmation sample.

## Appendix S3. Details associated w/ case study

The data used here were trail camera images classified via crowdsourcing from 91,276 24-h periods at 944 distinct locations across the state in 2017 between Julian days 150 and 320 (Figure S1). We defined sampling occasions as 24-hr periods, and reviewed all reported images reported as gray fox classifications (n =247 images) from 179 occasions at 127 locations; 90 other occasions at 55 distinct locations included putative but unconfirmed gray fox detections (Figure S1). Covariates for expected abundance are noted in Table S1 and were extracted from a circular buffer of 1 or 5 km radius surrounding the camera locations. Spatial smoothing was implemented via a 2-dimensional cubic spline (Guélat and Kéry 2018) across latitude and longitude with 20 knots placed across the state. The detection probability of an individual animal at different sites was modeled as varying in relation to whether the camera was placed on a maintained trail or not and as a quadratic function of the distance between the camera and the location the camera was targeting (as reported by volunteers), and false positive error probability was modeled as a logistic function of the proportion of cropland within a circular buffer with 5 km radius to account for what we expected to be increased prevalence of species confused with gray foxes (red fox, coyote) relative to foxes themselves. The prediction grain (a 2 × 2 km lattice) was chosen to approximate gray fox home range sizes in Wisconsin, which are believed to be slightly larger than the home ranges reported slightly further south (e.g., Haroldson and Fritzell 1984, Duell et al. 2017).

In Clare et al. (2019), we noted that perhaps the most useful predictor for classification error across a range of species was the degree of unanimity in the crowdsourced classifications (e.g., 100% of votes for gray fox = more likely to be gray fox than 50% votes). We do not use this term here to avoid the inelegance of having to define/impute values associated with crowdsourced agreement within sampling occasions in which the gray fox was not detected, although following the logic that greater confidence leads to lower probability of a false positive, imputing these values as 1.0 (i.e., 100% agreement) seems like a reasonable hack. Similar inelegancies associated with agreement arise for sampling occasions with multiple images; one way to model this that might be reasonable might be to define a covariate (or the false positive probability) as 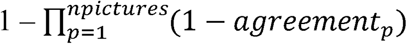. For example, in a case with 2 pictures in with 50 and 40 % agreement, the value of the operator = 0.7, in a case with 1 picture with 90% agreement, the value = 0.1, which seems to correctly imply a false positive is more likely in the first case because there are more images with less confidence. Of course, in situations with 0 pictures, the operator breaks down, and the value might need to be fixed at one.

Of the reviewed images, we confirmed 67% as correct classifications, with the rest either misclassifications of coyote (*Canis latrans*) or red fox (*Vulpes vulpes*). Once aggregated within sampling occasions, 60% (108) of the occasions consisted exclusively of true positives, and 40% (71) included exclusively false positives; no confirmed sampling intervals included a mixture of true and false positive observations. This might suggest some potential lack of independence between true and false positive outcomes, but we ignore that here because volunteers view images on the crowdsourcing platform at random, which makes it difficult to imagine that misperception has any serial structure or exhibits any other form of dependence. We note that both true and false positives were reported at 6 locations. Estimates are summarized in tables S2 and S3.

---

**Figure S1.**
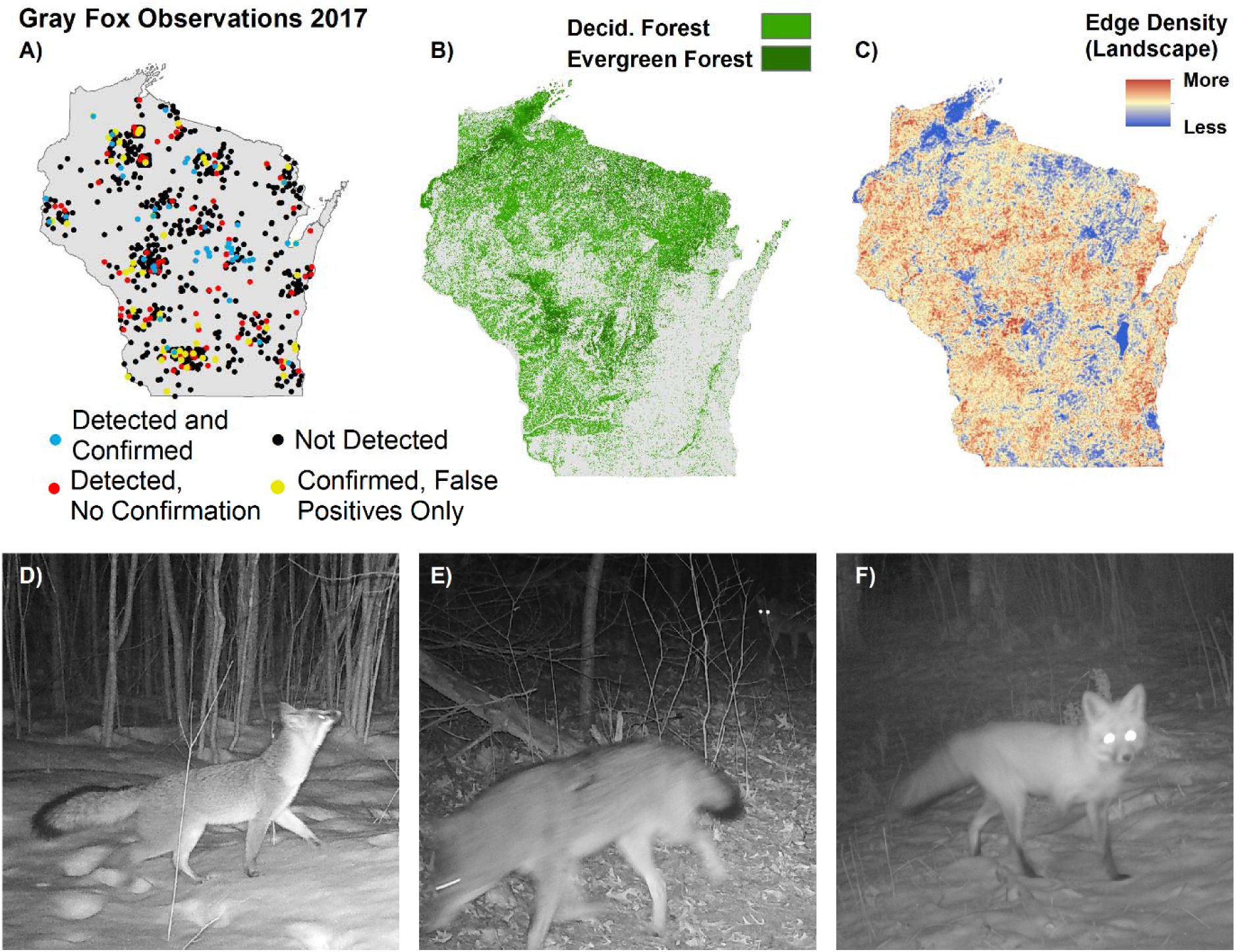
Location of sampling locations, confirmed observations, and unconfirmed detections used in the case study (A); covariates associated with model fitting (B and C). Reported gray fox (D) observations were commonly truly either coyote (E) or red fox (F).

**Table S1.**

| Name | Description | Source | Real Parameter |
| --- | --- | --- | --- |
| Evergreen | % Evergreen Forest in 1 km radius buffer surrounding camera location | NLCD 2011<sup>A</sup> | $\lambda$ |
| Snowdepth | Mean Snowdepth in 1 km radius buffer surrounding camera location between 2010 and 2017 | SNODAS<sup>B</sup> | $\lambda$ |
| Edge Density | (Landscape) edge density within 1 km radius buffer surrounding camera location | NLCD 2011 | $\lambda$ |
| Cropland | % Cropland in 1 km radius buffer surrounding camera location | NLCD 2011 | $\lambda, s_0$ |
| Grassland | % Grassland in 1 km radius buffer surrounding camera location | NLCD 2011 | $\lambda$ |
| Deciduous | % Deciduous forest in 1 km radius buffer surrounding camera location | NLCD 2011 | $\lambda$ |
| Trail | Binary; camera placed on maintained trail (vs. game trail or non-trail); | | $r$ |
| Distance | Distance between camera and target location (m); modeled as quadratic | | $r$ |

<sup>A</sup>Homer, C., Dewitz, J., Yang, L., Jin, S., Danielson, P., Xian, G., Coulston, J., Herold, N., Wickham, J. and Megown, K., 2015. Completion of the 2011 National Land Cover Database for the conterminous United States—representing a decade of land cover change information. *Photogrammetric Engineering & Remote Sensing* 81:345-354.
<sup>B</sup>Barrett, A. 2003. National Operational Hydrologic Remote Sensing Center SNOw Data Assimilation System (SNODAS) Products at NSIDC. NSIDC Special Report 11. Boulder, CO, USA: National Snow and Ice Data Center. Digital media.

**Table S2.**
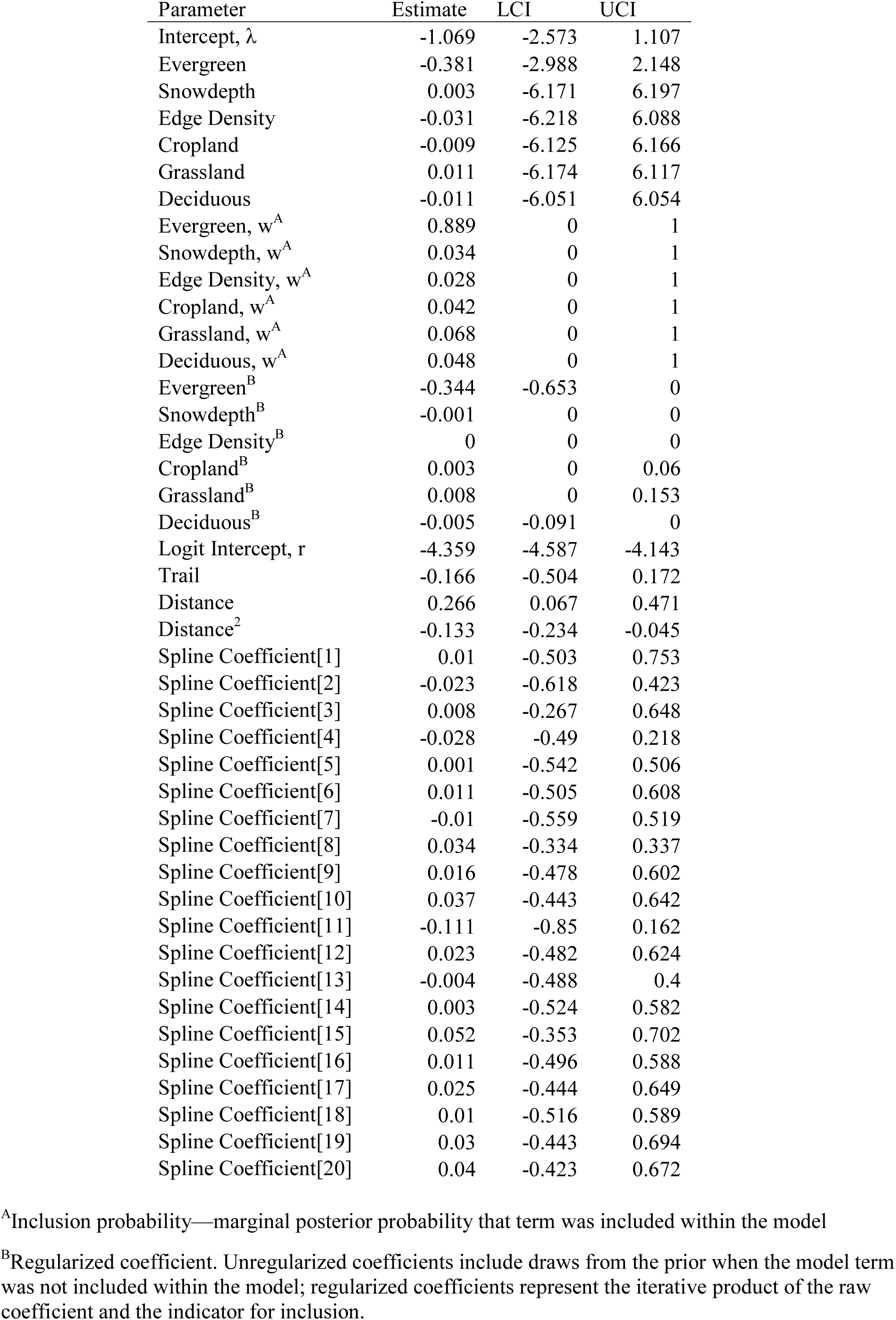
Parameter estimates, variable inclusion probabilities (*w*), and 95% credible intervals (LCI, UCI) from a RN model for gray fox relative abundance ignoring false positive error.

| Parameter | Estimate | LCI | UCI |
| --- | --- | --- | --- |
| Intercept, $\lambda$ | -1.069 | -2.573 | 1.107 |
| Evergreen | -0.381 | -2.988 | 2.148 |
| Snowdepth | 0.003 | -6.171 | 6.197 |
| Edge Density | -0.031 | -6.218 | 6.088 |
| Cropland | -0.009 | -6.125 | 6.166 |
| Grassland | 0.011 | -6.174 | 6.117 |
| Deciduous | -0.011 | -6.051 | 6.054 |
| Evergreen, $w^A$ | 0.889 | 0 | 1 |
| Snowdepth, $w^A$ | 0.034 | 0 | 1 |
| Edge Density, $w^A$ | 0.028 | 0 | 1 |
| Cropland, $w^A$ | 0.042 | 0 | 1 |
| Grassland, $w^A$ | 0.068 | 0 | 1 |
| Deciduous, $w^A$ | 0.048 | 0 | 1 |
| Evergreen<sup>B</sup> | -0.344 | -0.653 | 0 |
| Snowdepth<sup>B</sup> | -0.001 | 0 | 0 |
| Edge Density<sup>B</sup> | 0 | 0 | 0 |
| Cropland<sup>B</sup> | 0.003 | 0 | 0.06 |
| Grassland<sup>B</sup> | 0.008 | 0 | 0.153 |
| Deciduous<sup>B</sup> | -0.005 | -0.091 | 0 |
| Logit Intercept, $r$ | -4.359 | -4.587 | -4.143 |
| Trail | -0.166 | -0.504 | 0.172 |
| Distance | 0.266 | 0.067 | 0.471 |
| Distance<sup>2</sup> | -0.133 | -0.234 | -0.045 |
| Spline Coefficient[1] | 0.01 | -0.503 | 0.753 |
| Spline Coefficient[2] | -0.023 | -0.618 | 0.423 |
| Spline Coefficient[3] | 0.008 | -0.267 | 0.648 |
| Spline Coefficient[4] | -0.028 | -0.49 | 0.218 |
| Spline Coefficient[5] | 0.001 | -0.542 | 0.506 |
| Spline Coefficient[6] | 0.011 | -0.505 | 0.608 |
| Spline Coefficient[7] | -0.01 | -0.559 | 0.519 |
| Spline Coefficient[8] | 0.034 | -0.334 | 0.337 |
| Spline Coefficient[9] | 0.016 | -0.478 | 0.602 |
| Spline Coefficient[10] | 0.037 | -0.443 | 0.642 |
| Spline Coefficient[11] | -0.111 | -0.85 | 0.162 |
| Spline Coefficient[12] | 0.023 | -0.482 | 0.624 |
| Spline Coefficient[13] | -0.004 | -0.488 | 0.4 |
| Spline Coefficient[14] | 0.003 | -0.524 | 0.582 |
| Spline Coefficient[15] | 0.052 | -0.353 | 0.702 |
| Spline Coefficient[16] | 0.011 | -0.496 | 0.588 |
| Spline Coefficient[17] | 0.025 | -0.444 | 0.649 |
| Spline Coefficient[18] | 0.01 | -0.516 | 0.589 |
| Spline Coefficient[19] | 0.03 | -0.443 | 0.694 |
| Spline Coefficient[20] | 0.04 | -0.423 | 0.672 |

**Table S3.**
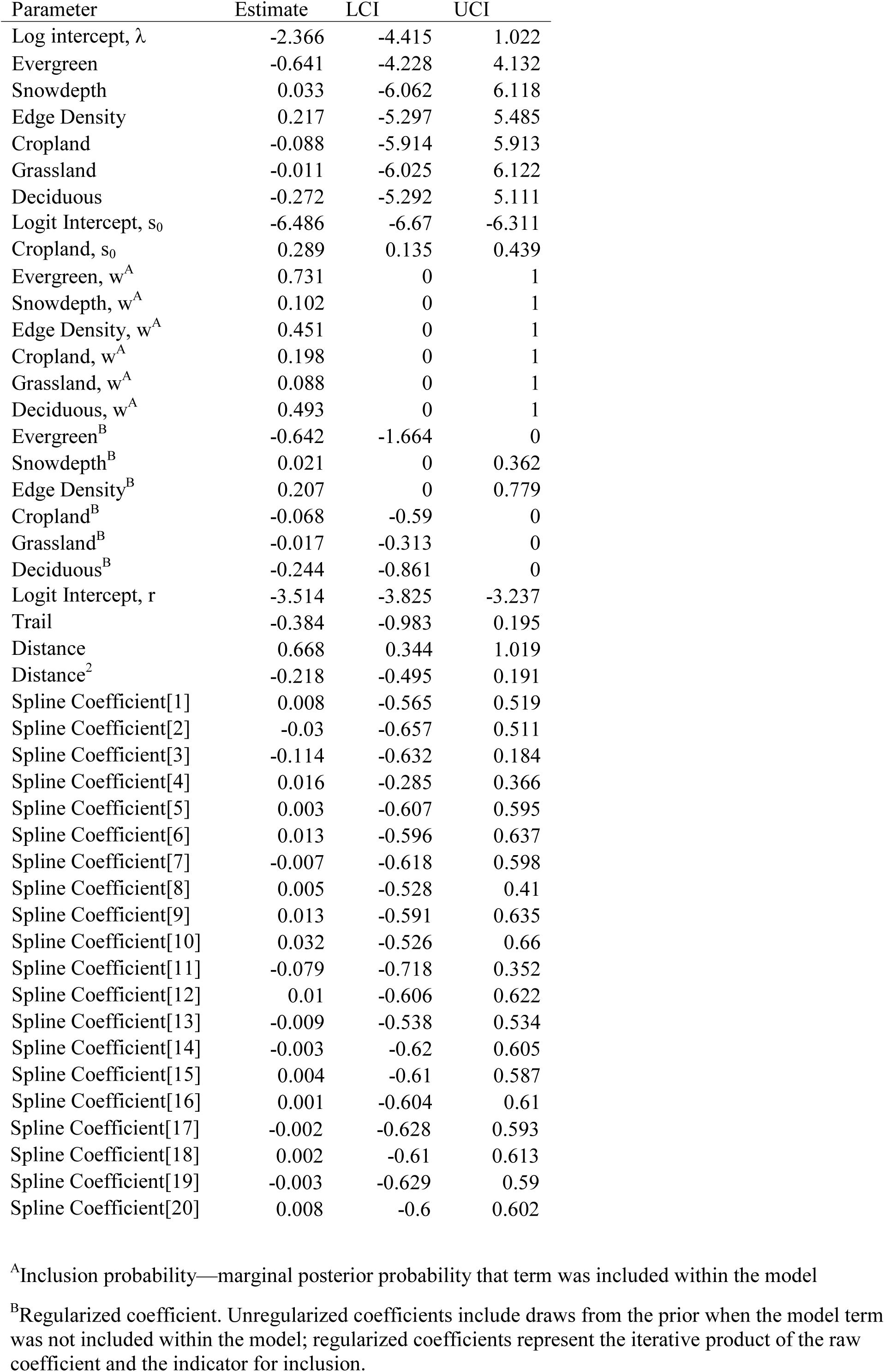
Parameter estimates, variable inclusion probabilities (*w*), and 95% credible intervals (LCI, UCI) from a RN model for gray fox relative abundance accounting for false positive error.

| Parameter | Estimate | LCI | UCI |
| --- | --- | --- | --- |
| Log intercept, $\lambda$ | -2.366 | -4.415 | 1.022 |
| Evergreen | -0.641 | -4.228 | 4.132 |
| Snowdepth | 0.033 | -6.062 | 6.118 |
| Edge Density | 0.217 | -5.297 | 5.485 |
| Cropland | -0.088 | -5.914 | 5.913 |
| Grassland | -0.011 | -6.025 | 6.122 |
| Deciduous | -0.272 | -5.292 | 5.111 |
| Logit Intercept, $s_0$ | -6.486 | -6.67 | -6.311 |
| Cropland, $s_0$ | 0.289 | 0.135 | 0.439 |
| Evergreen, $w^A$ | 0.731 | 0 | 1 |
| Snowdepth, $w^A$ | 0.102 | 0 | 1 |
| Edge Density, $w^A$ | 0.451 | 0 | 1 |
| Cropland, $w^A$ | 0.198 | 0 | 1 |
| Grassland, $w^A$ | 0.088 | 0 | 1 |
| Deciduous, $w^A$ | 0.493 | 0 | 1 |
| Evergreen<sup>B</sup> | -0.642 | -1.664 | 0 |
| Snowdepth<sup>B</sup> | 0.021 | 0 | 0.362 |
| Edge Density<sup>B</sup> | 0.207 | 0 | 0.779 |
| Cropland<sup>B</sup> | -0.068 | -0.59 | 0 |
| Grassland<sup>B</sup> | -0.017 | -0.313 | 0 |
| Deciduous<sup>B</sup> | -0.244 | -0.861 | 0 |
| Logit Intercept, $r$ | -3.514 | -3.825 | -3.237 |
| Trail | -0.384 | -0.983 | 0.195 |
| Distance | 0.668 | 0.344 | 1.019 |
| Distance<sup>2</sup> | -0.218 | -0.495 | 0.191 |
| Spline Coefficient[1] | 0.008 | -0.565 | 0.519 |
| Spline Coefficient[2] | -0.03 | -0.657 | 0.511 |
| Spline Coefficient[3] | -0.114 | -0.632 | 0.184 |
| Spline Coefficient[4] | 0.016 | -0.285 | 0.366 |
| Spline Coefficient[5] | 0.003 | -0.607 | 0.595 |
| Spline Coefficient[6] | 0.013 | -0.596 | 0.637 |
| Spline Coefficient[7] | -0.007 | -0.618 | 0.598 |
| Spline Coefficient[8] | 0.005 | -0.528 | 0.41 |
| Spline Coefficient[9] | 0.013 | -0.591 | 0.635 |
| Spline Coefficient[10] | 0.032 | -0.526 | 0.66 |
| Spline Coefficient[11] | -0.079 | -0.718 | 0.352 |
| Spline Coefficient[12] | 0.01 | -0.606 | 0.622 |
| Spline Coefficient[13] | -0.009 | -0.538 | 0.534 |
| Spline Coefficient[14] | -0.003 | -0.62 | 0.605 |
| Spline Coefficient[15] | 0.004 | -0.61 | 0.587 |
| Spline Coefficient[16] | 0.001 | -0.604 | 0.61 |
| Spline Coefficient[17] | -0.002 | -0.628 | 0.593 |
| Spline Coefficient[18] | 0.002 | -0.61 | 0.613 |
| Spline Coefficient[19] | -0.003 | -0.629 | 0.59 |
| Spline Coefficient[20] | 0.008 | -0.6 | 0.602 |

<sup>A</sup>Inclusion probability—marginal posterior probability that term was included within the model
<sup>B</sup>Regularized coefficient. Unregularized coefficients include draws from the prior when the model term was not included within the model; regularized coefficients represent the iterative product of the raw coefficient and the indicator for inclusion.

